# Neural correlates of embodied and vibratory mechanisms associated with vocal emotion production

**DOI:** 10.1101/2024.05.14.594073

**Authors:** Garance Selosse, Didier Grandjean, Leonardo Ceravolo

## Abstract

Despite a large body of literature on the psychological and brain mechanisms of vocal emotion perception, less is known on expression and production mechanisms, especially the vibrations originating in the vocal cords and their role in emotional voice production. In the present study, we aimed to fill this gap. Participants had to produce angry, happy and neutral emotional vocalizations in different production conditions (‘normal’, ‘whisper’, ‘articulate’). An accelerometer recorded the vibrations on the throat, close to the vocal folds. Results showed effects of the Emotion factor with activations in the bilateral temporal voice areas, the inferior frontal gyri as well as motor and supplementary motor areas. The Production factor and its interaction with Emotion revealed significant effects in motor, somatosensory cortices, insula and inferior frontal cortex. Exploratory analysis of the brain correlates of emotional vocal tract vibrations specific to ‘normal’ voice production showed significant correlations with brain regions involved in interoceptive processing (insula, inferior frontal cortex, cerebellum). Our results highlight the crucial role of vibro-tactile body resonances in vocal emotion production that might play an important role for the interoceptive phenomena involved in the representation of our own emotions such as in emergent feelings related to emotional vocal production.

## Introduction

Oral communication is of crucial importance for social creatures such as us humans, involving not only the semantic meaning of words but also non-verbal cues. The term ‘emotional voice prosody’ encompasses changes in acoustic parameters that complement or even add new information to speech (Grandjean et al., 2006; Laukka et al., 2005; Risberg & Lubker, 1978; Scherer & Oshinsky, 1977). It refers, at the suprasegmental level, to the perceived melodic aspects of speech, which include variations in pitch and timing, as well as spectral fluctuations related to timbre and formants which are more related to segmental aspects of speech. These acoustic parameters are primarily influenced by the mechanisms involved in the production of vocalizations, including breathing, phonation and articulation, which in turn are modulated by emotions (Banse & Scherer, 1996). Consequently, emotional prosody conveys crucial information about the speaker’s affective state and plays a central role in regulating social interactions (Pell, & Kotz, 2011; Sauter & Eimer, 2010).

While there has been substantial research into the neural mechanisms underlying the perception of emotional prosody, the neural mechanisms involved in its *production* remain less understood and vastly understudied. In recent years, a slowly increasing number of neuroimaging studies have provided valuable insights into this complex process. Several studies of brain-damaged patients and healthy neuroimaging studies have shown the importance of the right frontal cortex in the production of emotional vocalizations (Dogil et al., 2002; Riecker et al., 2005; Ross & Monnot, 2008). Indeed, Wright et al. (2016) tested brain-damaged patients with impairments in the expression or/and recognition of affective prosody. They found that patients with selective impairments in the expression of affective prosody had right inferior frontal lesions. Other studies investigating healthy participants have shown that the inferior frontal gyrus was involved in both the perception and production of emotional prosody, illustrating its potential implication in a network in which one’s own voice is monitored and adjusted by sensory feedback and top-down processes (Aziz-Zadeh et al., 2010; Zheng et al., 2010).

Other structures were reported to be important for vocal emotion production. Firstly, we should mention the basal ganglia. It was reported to be involved in the volitional control of vocal affect (Laukka et al., 2011). More specifically, studies claimed its implication during the preparation phase of emotional prosody production and a functional segregation for emotional and sensorimotor/cognitive processes. Indeed, the bilateral ventral striatum is linked to limbic structures, temporal poles and anterior insula, and appears to be activated for the preparation of emotional prosody production (Pichon & Kell, 2013; Yelnik, 2008). As for the bilateral dorsal striatum, it is deeply connected with sensorimotor and associative cortices (Yelnik, 2008) and seems to be more associated with cognitive and motor aspects of the preparation (Pichon & Kell, 2013). Second, the superior temporal gyrus is claimed to have a role in processes such as the provision of articulatory maps (Aziz-Zadeh et al., 2010; Pichon & Kell, 2013), sensory motor integration (Hickok, 2009; Peschke et al., 2009), and phonological feedback processing (Dogil et al, 2022). Then, the anterior cingular cortex (ACC) is associated with the monitoring of vocal production (Jurgens, 2009; Hage, 2010; Frühholz et al., 2015). Both the ACC and the insula appear to be involved in regulating autonomic arousal when people produce aggressive vocalizations (Frühholz et al., 2015). The insula is also recruited for the execution of speech movements but not for articulatory planning (Dogil et al., 2022). It was also highlighted as a region involved in interoceptive awareness and emotional experience (Zaki et al., 2012). Yet another cerebral structure crucially involved in speech production is the cerebellum, as it was reported to be responsible for the fine control of motor speech, especially for temporal aspects (Manto et al., 2012). Moreover, a study investigating the production of emotional speech revealed significant activations in the cerebellar vermis associated with temporal variations in fundamental frequency (Pichon & Kell, 2013). Finally, the motor network including the supplementary motor area and the motor cortex is associated with the planning of articulatory movements (Dogil et al., 2022) and the control of the motor output (Alario et al., 2006; Frühholz et al., 2015). We can therefore appreciate the large network of cortical and subcortical brain regions involved in vocal emotion production.

In the present study, we were essentially interested in a so far neglected aspect of voice production, namely vocal cord vibrations concomitant to emotional speech production. We therefore focused on varying types of vocal emotions and prosody production conditions (whispering, articulating, and normal speech) that led to the selective vibration—normal speech—or absence of vibration—whispering, articulating—of the vocal cords. Such manipulation elicited distinct bodily resonances in our participants, thereby enhancing our understanding of the interplay between vocal production and interoceptive processes. Interoception is defined in the literature as the sense of the body’s internal physiological state and includes the perception of sensations such as heartbeat, breathing and visceral processes, allowing individuals to maintain awareness of their physical condition (Craig, 2002; Schulz, 2016). Insular activity is of particular interest on this topic due to its close ties with interoception and body awareness mechanisms. In fact, it was speculated that the insular cortex could potentially serve as a central area in the integration of somato-visceral sensations, subjective feelings, cognitive appraisals, and the consciousness of internal and external physiological signals (Adolfi et al., 2017; Casals-Gutierrez, & Abbey, 2020; Grandjean et al., 2008). Most of the studies investigating interoception used heartbeat perception tasks and they showed activation(s) in the insular cortex—but also in other structures such as the cingular cortex, the somatosensory cortex, the supplementary motor area (SMA) and the prefrontal cortex (Engelen et al., 2023; Gibson, 2019; Khalsa et al., 2018).

In our study, we took the path of vocal emotion production and vocal cord vibrations as a type of potential interoceptive signal, assuming similar results as the ones mentioned above would be observed especially in the insula and somato-sensory regions. We therefore investigated the mechanisms involved in the production of emotional prosody, following on from the work previously carried out by others in the recent past. Our participants had to vocally produce according to different production conditions such as normal production, articulation and whispering, to explore the different nuances linked to emotional—angry, neutral, happy— production, considering the presence or absence of vocal cords vibrations in our production conditions. These three production conditions therefore also differed in terms of one’s own auditory and sensory feedback—presence or absence of vocal cords vibrations. Furthermore, we chose to consider one positive and one negative emotion with happiness and anger—high in arousal, in addition to a neutral condition. We thus hypothesized: i) normal—compared to whispering and articulating—emotional compared to neutral productions to enhance activity previously observed in neuroimaging studies on emotional speech production such as the IFG (Aziz-Zadeh et al., 2010; Holland et al., 2011), the basal ganglia (Pichon & Kell, 2013; Yelnik, 2008) and the superior temporal cortex (Aziz-Zadeh et al., 2010; Dogil et al., 2022; Pichon & Kell, 2013); ii) due to the presence of concomitant vocal cord vibrations for normal emotional vs. neutral speech production—compared to whispering and articulating—and their potential to involve somato-sensory and interoceptive brain areas, we also expected enhanced activity in supplementary motor area and somatosensory regions (Alario et al., 2006), as well as in the insula (Dogil et al., 2022; Zaki et al., 2012). These effects were expected to be stronger for emotional than neutral produced voices and in a similar fashion for angry and happy voices. iii) correlates of voice vibrations—generated in the ‘normal’ speech production condition only, especially for produced vocal emotion, in the superior temporal cortex (Aziz-Zadeh et al., 2010; Hickok, 2009; Dogil et al, 2022), somato-sensory brain areas (Adolfi et al., 2017; Casals-Gutierrez, & Abbey, 2020) and the cerebellum (Manto et al., 2012; Pichon & Kell, 2013).

## Methods

### Participants

For the main voice production task, twenty-five healthy volunteers were recruited amongst psychology students from the University of Geneva (twelve females, thirteen males; *M_Age_*=22.88; *SD_Age_*=3.20). Sample size was decided *a priori* upon using a power analysis adjusted to the design of our study with an effect size of 0.5, a power of 0.8 and an alpha of 0.05, resulting in a required sample size of N=25 (G*Power 3; Faul et al., 2007).

For the temporal voice areas functional localizer task, a sample of 117 participants was gathered (55 male, 62 female, *M_Age_*=24.82 years, *SD_Age_*=5.45).

For both samples, all participants were at least 18 years old, naïve to the tasks, reported normal hearing and no neurologic or psychiatric history. Participants gave informed and written consent for their participation in the experiment. The study was approved by the local ethics committee in accordance with ethical and data security guidelines of the University of Geneva and conducted according to the declaration of Helsinki.

### Stimulus material: voice production task

While lying in the MRI scanner, participants were presented with written pseudowords displayed on a grey screen with Matlab (The Mathworks Inc., Natick, MA, USA), using the Psychophysics Toolbox 3 extensions (Brainard, 1997; Pelli, 1997; Kleiner et al., 2007). The pseudowords were selected to avoid any semantic meaning that might interfere with the production of emotional voices. Four different pseudowords (‘belam’, ‘nolan’, ‘goster’, and ‘semina’) were chosen, taken from the highly validated Geneva Multimodal Emotion Portrayal corpus (Bänziger & Scherer 2010).

### Stimulus material: temporal voice areas localizer task

Auditory stimuli consisted of sounds from a variety of sources. Vocal stimuli were obtained from 47 speakers: 7 babies, 12 adults, 23 children and 5 older adults. Stimuli included 20 blocks of vocal and 20 blocks of non-vocal auditory stimuli. Vocal stimuli within a block could be either speech (33%: words, non-words, foreign language) or non-speech (67%: laughs, sighs, various onomatopoeia). Non-vocal stimuli consisted of natural sounds (14%: wind, streams), animals (29%: cries, gallops), the human environment (37%: cars, telephones, airplanes) or musical instruments (20%: bells, harp, instrumental orchestra). The paradigm, design and stimuli were obtained through the Voice Neurocognition Laboratory website (http://vnl.psy.gla.ac.uk/resources.php). Stimuli were presented at an intensity that was kept constant throughout the experiment (70 dB sound-pressure level). Participants were instructed to actively listen to the sounds but no other task was required. The silent inter-block interval was 8sec long for a total task duration of 11min.

### Task procedure: voice production task

During the experiment, six runs were performed expressing either anger, happiness or neutral emotions (two runs each). Each block contained one mini-block per production condition to avoid too much switching between these conditions that required concentration. Each of these mini-blocks consisted of eighteen voice outputs, for a subtotal of 54 outputs per block and a total of 324 outputs per participant (3 production conditions x 18 voice outputs x 6 blocks). For one emotion, there were in total 36 trials per production type, for a grand total of 108 trials per emotion across production types. One emotion was always represented solely in one full run to avoid confusions and to once again have a less complicated instruction-switching task. The order of the runs was pseudo-randomized and counterbalanced between participants but the same emotion could not appear twice in a row. In addition to the emotion instruction, participants had the following production instructions (mini-blocks) within the runs: pronounce in a normal way (‘Normal’ condition), whisper (‘Whisper’ condition) or articulate only (‘Articulate’ condition). Each condition was explained in detail and rehearsed before each participant entered the MRI scanner. These conditions allow for body resonances and auditory feedback control—of no-interest—through the presence/absence of vibrations and/or phonation, respectively. They had to produce the pseudowords one by one, in one go, and in the most natural way in a specific production condition and expressing an instructed emotion in a period of up to 3 seconds (Figure 1), during which the scanner was silent. Between each trial, a blank screen of 5 seconds was displayed. This time interval served as a window to acquire fMRI volumes in a task-triggered, sparse-sampling fashion.

**Figure 1.**
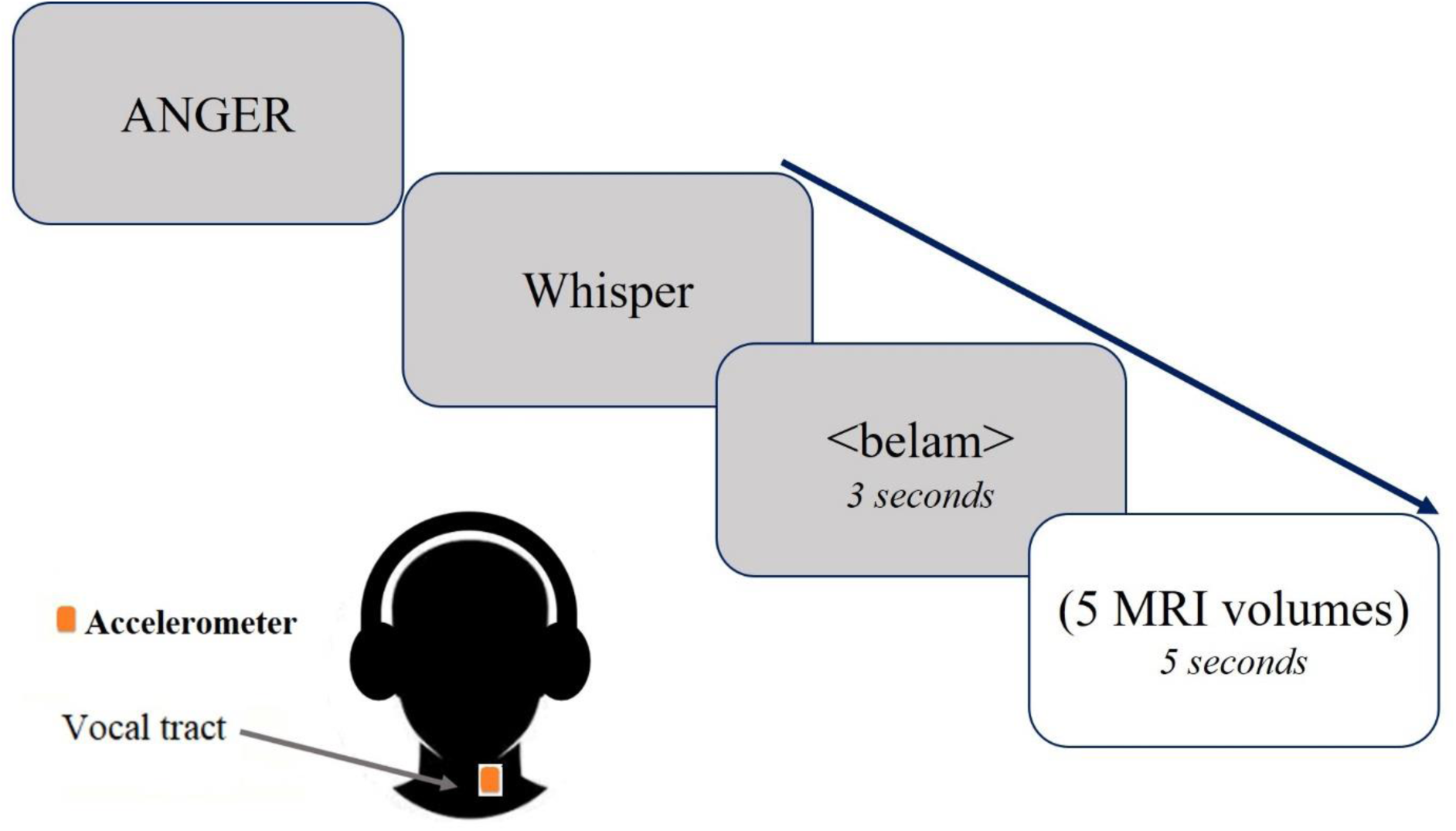
Experimental design. Participants were instructed to produce pseudo-words expressing either anger, happiness or neutral emotions (two runs of each emotion), following a Production instruction (‘Normal’, ‘Whisper’ or ‘Articulate’). Each run consisted of one mini-block per production condition. They had to produce the pseudowords one by one, in one go, and in the most natural way possible within a period of three seconds. A blank screen of five seconds was displayed between each trial.

### fMRI data acquisition and analysis

All functional imaging data were recorded on a 3-T Siemens Trio System (Siemens, Erlangen, Germany) scanner equipped with a 32-channel antenna. All information and details concerning MRI data acquisition and analysis are reported in the supplementary material for the production task and the TVA task.

## Results

The task was a production task of pseudowords (‘belam’, ‘nolan’, ‘goster’, and ‘semina’) following different production instructions (‘normal’, ‘whisper’, and ‘articulate’) and expressing three emotions (anger, happiness and neutral). The factors of interest were therefore the Emotion, the Production and the interaction between these two factors. Twenty-five (12 females) participants were included in the final analysis of the present study for the wholebrain and ROI analyses of the vocal emotion production task while vocal tract vibrations correlates included sixteen participants, due to data contamination. These data are presented as exploratory for this reason. More details on the task and paradigm can be found in the Methods section.

### Wholebrain results

#### Main effects of Emotion and Production factors and their interaction

Whole-brain results for the Emotion factor showed that enhanced activations for emotional (angry and happy) compared to neutral voices were significant especially in the TVA such as the superior temporal cortices and inferior frontal cortices, bilaterally, as well as in the insula, the planum temporale, SMA, motor and premotor areas (Figure 2, Supplementary Table 1). Results also showed significantly increased activity for both angry > neutral voices and happy > neutral voices, more details are available in Figure S2 and S3.

**Figure 2.**
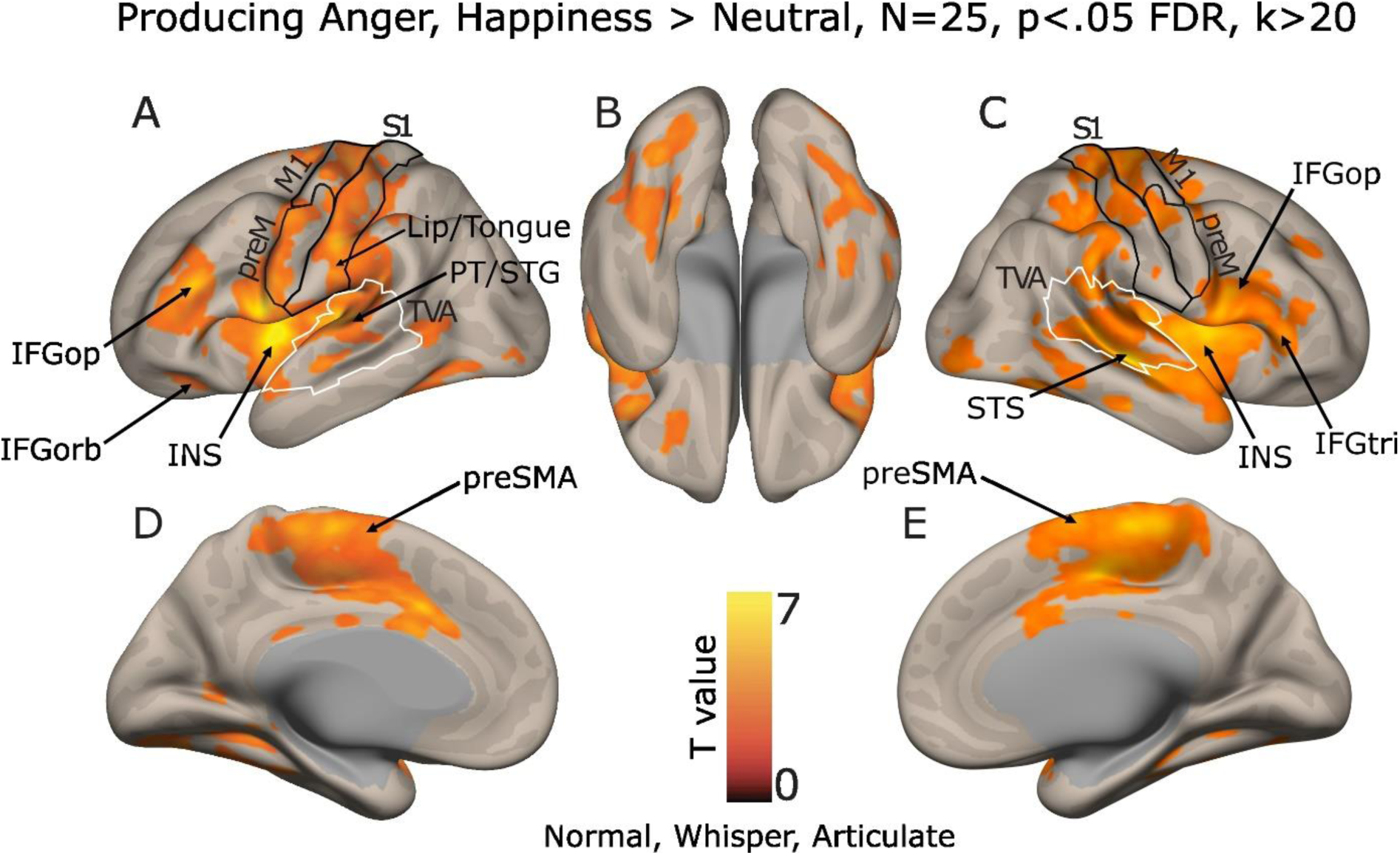
Brain measures for the production of emotional (happy and angry) compared to neutral voices, corrected for multiple comparisons (wholebrain voxel-wise *p* < .05 FDR, k > 20 voxels). The colorbar represents the statistical T value. IFG: inferior frontal gyrus, IFGop: pars opercularis, IFGorb: pars orbitalis, IFGtri: pars triangularis INS: insula, M1: primary motor cortex, preM: premotor area, PT: planum temporale, S1: primary somatosensory cortex, preSMA: anterior portion of the supplementary motor area, STG: superior temporal gyrus, STS: superior temporal sulcus, TVA: Temporal Voice Areas delineated by white line shapes.

Brain activity for angry compared to happy voices was observed in the TVA such as in the middle superior temporal gyrus, as well as in the *pars opercularis* of the inferior frontal gyrus, the SMA, motor and premotor areas as well as in the ventromedial prefrontal cortex (Figure 3). More details are available in Supplementary Table 2.

**Figure 3.**
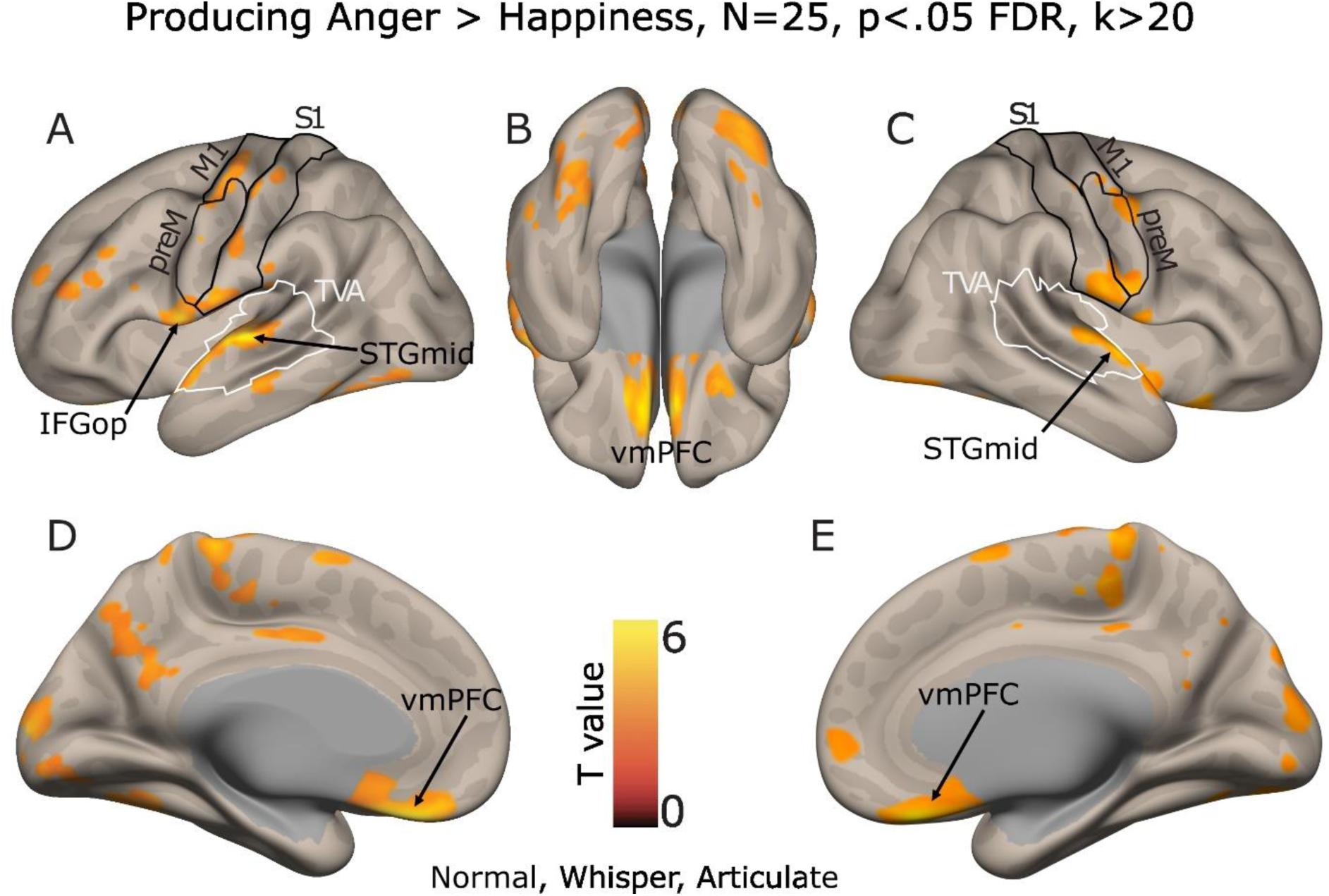
Brain measures for the production of angry compared to happy voices, corrected for multiple comparisons (wholebrain voxel-wise *p* < .05 FDR, k > 20 voxels). The colorbar represents the statistical T value. IFG: inferior frontal gyrus, M1: primary motor cortex, op: pars opercularis, preM: premotor area, S1: primary somatosensory cortex, preSMA: anterior portion of the supplementary motor area, STGmid: middle superior temporal gyrus, TVA: Temporal Voice Areas delineated by white outlines, vmPFC: ventromedial prefrontal cortex.

Brain activity for happy compared to angry voices showed the involvement of the superior temporal cortex and the inferior frontal gyrus (Figure 4). Enhanced activity was also observed in the supramarginal gyrus, the insula, the middle temporal gyrus, the parahippocampal gyrus, the ventromedial prefrontal cortex, the primary somatosensory cortex, the premotor and motor areas. More details are available in Supplementary Table 3.

**Figure 4.**
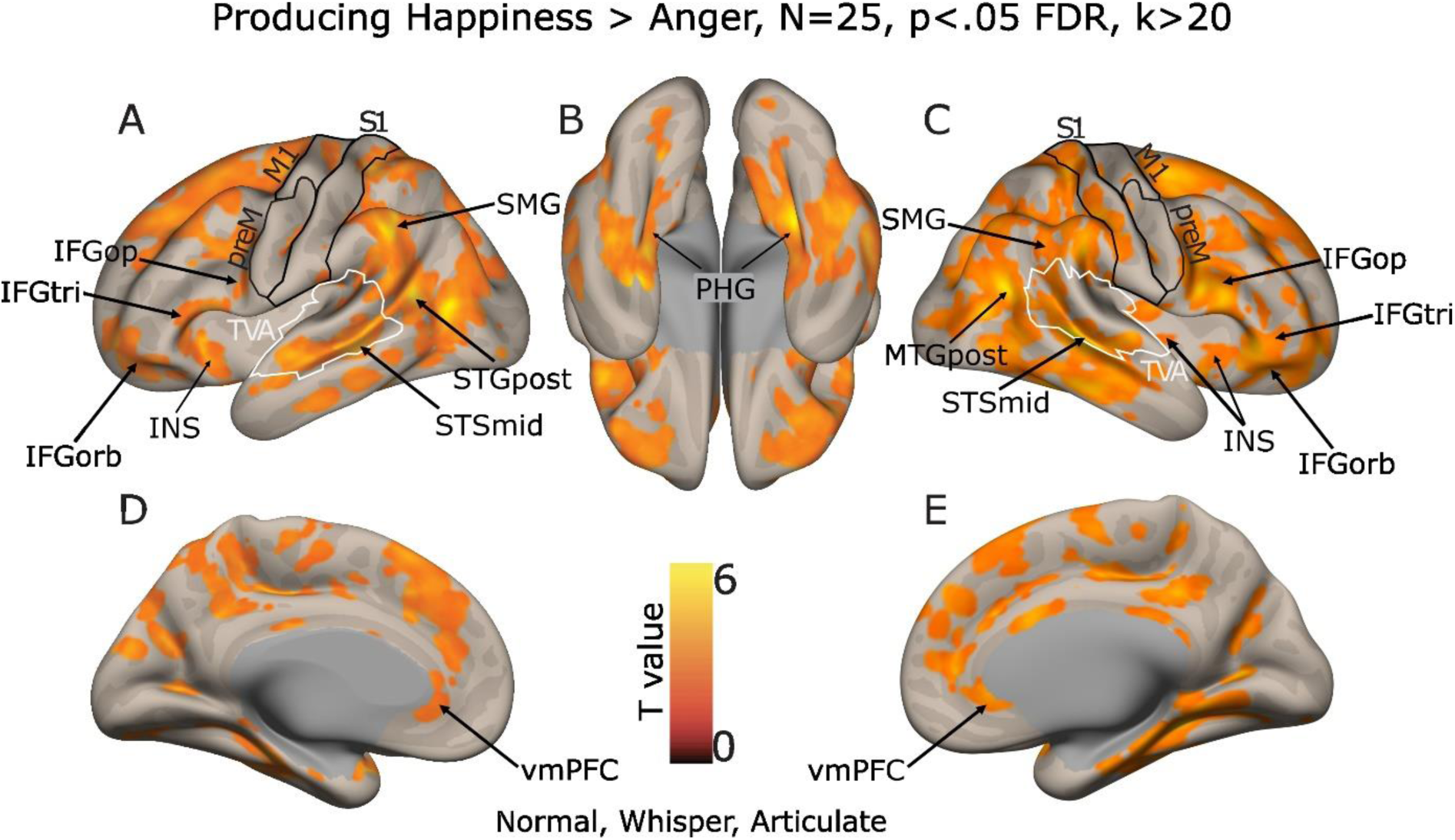
Brain measures for the production of happy compared to angry voices, corrected for multiple comparisons (wholebrain voxel-wise *p* < .05 FDR, k > 20 voxels). The colorbar represents the statistical T value. IFG: inferior frontal gyrus, INS: insula, M1: primary motor cortex, MTG: middle temporal gyrus, op: pars opercularis, orb: pars orbitalis, preM: premotor area, PHG: parahippocampal gyrus, S1: primary somatosensory cortex, preSMA: anterior portion of the supplementary motor area, SMG: supramarginal gyrus, STG: superior temporal gyrus, STS: superior temporal sulcus, tri: pars triangularis, TVA: Temporal Voice Areas delineated by white line shapes, vmPFC: ventromedial prefrontal cortex.

Wholebrain analysis for the Production factor did not show any effects, neither did the interaction between Emotion and Production. As planned, we then conducted a mask-inclusive analysis on our regions-of-interest (N=28’332 voxels, see Methods) based on the [anger, happiness > neutral voice production] contrast, including the TVA, the IFG, somatosensory and motor cortices, the insula, to further investigate the Production factor (Figure 5). Neural activations were observed in all of these regions. Moreover, statistics were computed between the conditions of the Production factor (Figure 5F). Significant results were found in the left middle and posterior insula, the left primary somatosensory cortex, the right middle STS, and bilaterally in the primary motor cortex. Specifically, there were significant contrasts between ‘normal’ and ‘whisper’ and ‘articulate’ conditions, respectively, in the left middle insula (Figure 5A and F); between ‘normal’ and ‘articulate’ conditions in all regions, with a tendency toward significance only for the left posterior insula; between ‘normal’ and both ‘articulate’ and ‘whisper’ conditions in the right middle STS and left primary motor cortex, with a tendency toward significance for the right hemisphere. More details are available in Supplementary Table 4.

**Figure 5.**
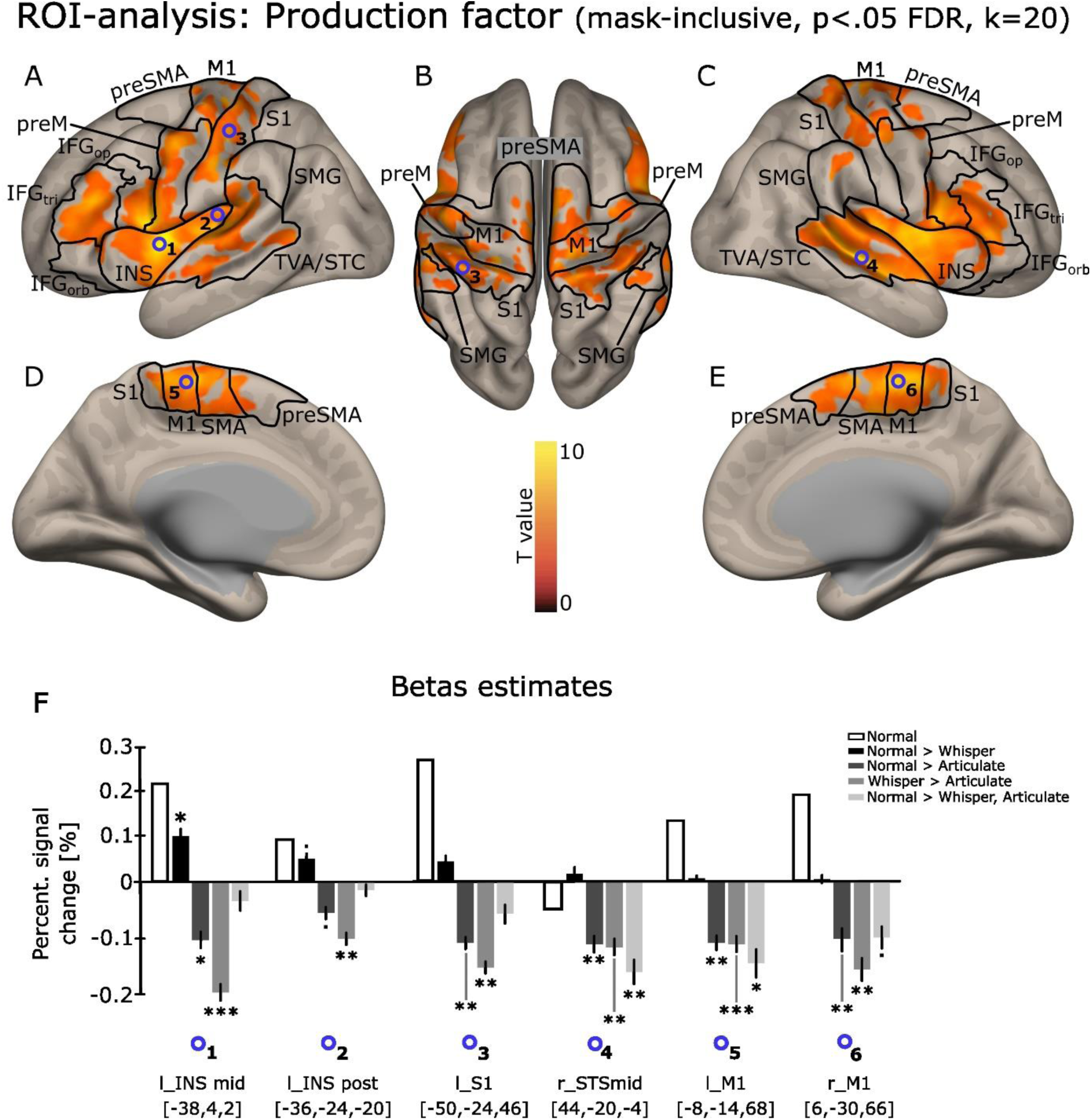
Brain activity in our regions of interest (ROI-analysis, mask-inclusive) for the Production factor, corrected for multiple comparisons (wholebrain voxel-wise *p* < .05 FDR, k > 20 voxels) and based on the [Anger, Happiness > Neutral] contrast (A-E). The colorbar represents the statistical T value. (F) Betas were extracted (blue circles represent significant activity peaks) using the group-level activity peaks within the ROIs: the nearest local maxima within a radius of 3mm was searched for each participant and then a cube of 27 contiguous voxels was used around this selected peak, with a singular value decomposition among these 27 voxels to select the ones that explained at least 90% of the variance. Statistics were then computed using repeated measures ANOVAs with factors Emotion and Production type. Histograms represent the percentage of signal change. IFG: inferior frontal gyrus, INS: insula, M1: primary motor cortex, SMA: supplementary motor area, preM: premotor area, S1: primary somatosensory cortex, preSMA: anterior portion of the supplementary motor area, SMG: supramarginal gyrus, STC: superior temporal cortex, TVA: Temporal Voice Areas, tri: *pars triangularis*, op: *pars opercularis*, orb: *pars orbitalis*. ·.05<p<.1; *p<.05; **p<.01; ***p<.001.

Among these regions of interest, only the left mid insula and right mid STS showed a significant interaction between Emotion and Production factors (left mid insula: *F*(4,96)= 4.30, *p*<.01; right mid STS: *F*(4,96)= 5.29, *p*<.001). We computed contrasts for these interactions to test our hypotheses and only one planned contrast was significant, in the right mid STS ([Emotion: Happiness > Neutral] x [Production: Normal > Whisper, Articulate]; *t*(24)=-2.54, *p*<.05). Upon closer inspection, the interaction appears to be driven by ‘neutral’ emotion and ‘articulate’ production conditions as shown in Supplementary Table 5 and 6.

#### Correlates of vocal tract vibrations during normal voice production

Here we present the exploratory results of the brain correlates of emotional vocal tract vibrations specific to ‘normal’ voice production (Production factor for normal voice as well as the Emotion*Production interaction, Figure 5)—‘whisper’ and ‘articulate’ conditions did not have any associated vibrations. Noteworthy is the fact that here we are not looking at enhanced brain activity according to conditions onsets, but rather to enhanced activity correlating with increased and decreased averaged vocal tract vibrations, across trials of a similar condition. In order to show all interaction contrasts of interest, the statistical thresholds used here are not as conservative as those in the results above—even though some, but not all of the following results survive higher statistical thresholding (see Supplementary Table 7). This is mainly due to technical problems during data collection of the vibrations and irreversible signal noise— namely, the extremely high sensitivity of the accelerometers that unwillingly collected scanner vibrations in a number of our participants. For these analyses, sample size therefore had to be reduced to 16 individuals. We present in this section the vocal tract vibratory correlates for the interaction between the Emotion and Production factors while the exploratory results for the Emotion factor as well as the Production factor can be found in Figure S4, S5, S6, respectively.

Results specific to normal voice production vibrations, across emotions, were found in the left dorsolateral prefrontal cortex (DLPFC), posterior STG (Figure 6A) and caudate nucleus as well as in the right mid STS (Figure 6B). Concerning interaction contrasts, we found correlates of vocal tract vibrations for both angry and happy compared to neutral ‘normal’ vocal productions in the medial orbitofrontal cortex (OFC), left cerebellum (Crus I region; Figure 6C) as well as in the right posterior middle temporal gyrus (MTG) and anterior inferior temporal sulcus (ITS; Figure 5D). For the angry versus happy ‘normal’ voice production contrast, vocal tract vibrations correlates were observed again in the medial OFC, left anterior MTG and in the left cerebellum, lobule VI (Figure 6E) as well as in the left posterior STG and anterior and posterior MTG (Figure 6F). For the inverse contrast finally—happy versus angry ‘normal’ voice production, vibrations correlates were observed in the left anterior insula, right fusiform cortex (FFC; Figure 5G) and left IFG *pars opercularis* (Figure 5H). Several temporal lobe correlates interestingly fall within the bounds of the TVA (black outline in Figure 6A,B,F).

**Figure 6.**
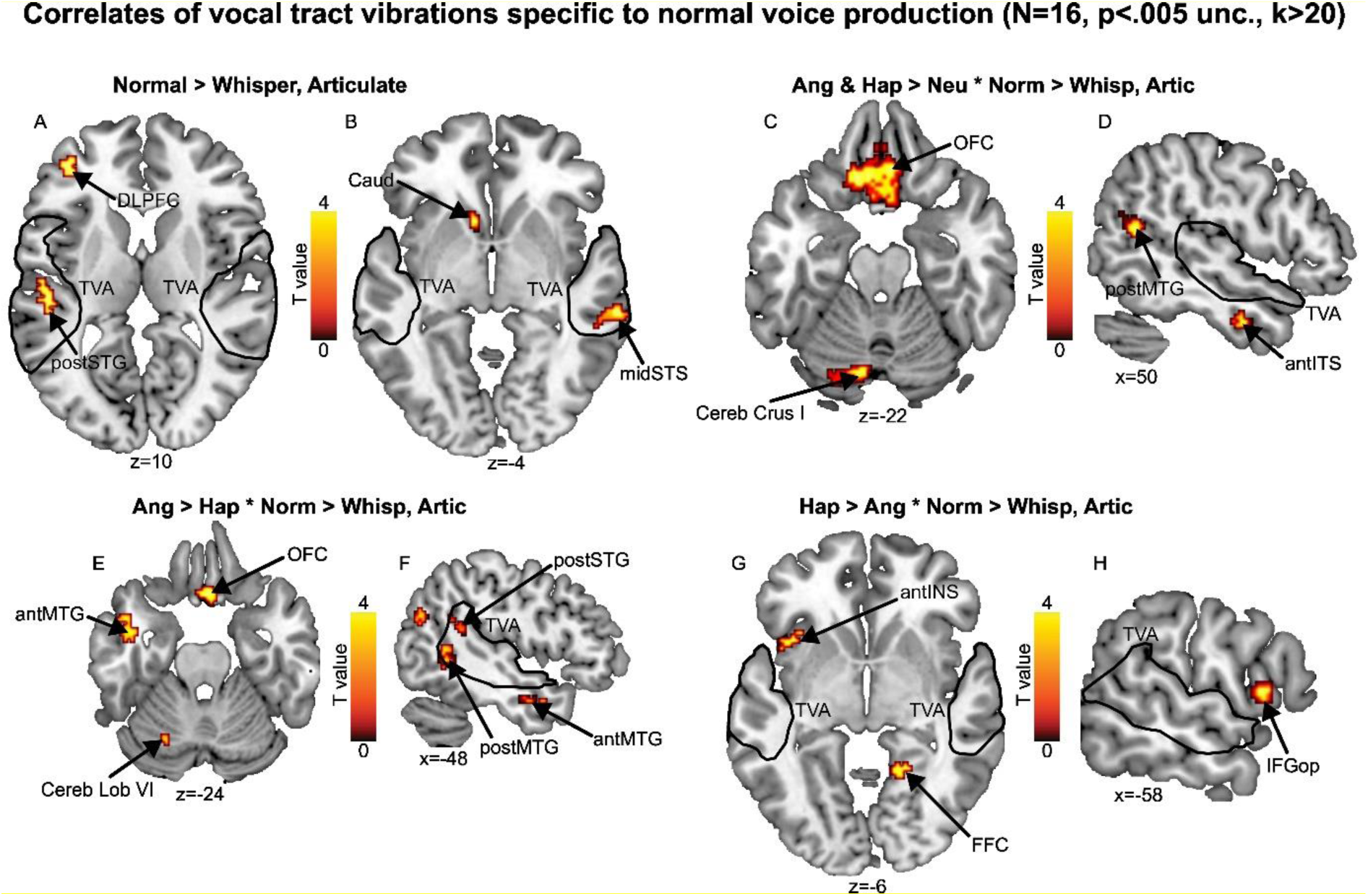
Brain correlates of vocal tract vibrations. A,B Brain correlates for the production factor. C,D Vibrations of normally produced angry and happy compared to neutral voices. E,F Vibrations of angry compared to happy normal voice production. G,H Vibrations of happy compared to angry normal production voices. Statistical thresholding was set to an uncorrected wholebrain voxel-wise *p* < .005, k > 20 voxels. The colorbars represent the statistical T value. DLPFC: dorsolateral prefrontal cortex, STG: superior temporal gyrus, Caud: caudate nucleus, STS: superior temporal sulcus, OFC: orbitotronfal cortex, Cereb: cerebellum, MTG: middle temporal gyrus, ITS: inferior temporal sulcus, Cereb Lob: cerebellum lobule, INS: insula, FFC: fusiform cortex, IFGop: inferior frontal cortex *pars opercularis*; ‘ant’ prefix: anterior, ‘mid’ prefix: mid portion, ‘post’ prefix: posterior, TVA: temporal voice areas delineated by black outlines.

## Discussion

While there has been considerable research on the neural mechanisms underlying the perception and the processing of emotional prosody, the neural mechanisms involved in vocal emotion production remain understudied and therefore significantly less understood. In the present study, we aimed to investigate the production of emotional prosody using different production conditions (normal speech, whispering, articulating) that would lead to the selective activation—or absence thereof—of vocal cords vibrations and/or phonation. Our main hypothesis was that normal emotional production would increase activity in the IFG, basal ganglia, supplementary motor and somatosensory area, superior temporal cortex and insula. This effect was expected to be stronger for emotional than for neutral voices as well as for angry compared to happy voices. Vocal emotion production triggered wholebrain activity consistent with previous literature and with our hypothesis, while ROI-analysis highlighted enhanced activity for specific production conditions in the insula, motor and somatosensory brain areas. The exploration of brain correlates of vocal tract vibrations during normal production suggests a contribution of sensory processing mechanisms in the production of vocal emotions.

Wholebrain results for the Emotion factor globally showed activations in voice-sensitive brain areas such as the superior temporal gyrus and sulcus and the inferior frontal cortex. This is consistent with our hypotheses on the voice production task and is in line with existing literature. In fact, the implication of these brain regions in processes such as the provision of articulatory maps (Aziz-Zadeh et al., 2010; Pichon & Kell, 2013), sensory motor integration (Hickok, 2009; Peschke et al., 2009), and phonological feedback processing (Dogil et al, 2022) by the superior temporal cortex, as well as one’s own voice monitoring by sensory feedback and top-down processes by the frontal gyrus (Aziz-Zadeh et al., 2010; Zheng et al., 2010). These activations confirmed—and replicate—the existence of a common network between voice *perception* and *production* (Scott, 2022; Parkinson et al., 2012). Moreover, brain activations in the motor network, i.e., the SMA, motor and premotor areas were also observed. This network was previously highlighted in vocal production at several levels such as the planning of articulatory movements (Dogil et al., 2022), the control of the motor output (Alario et al., 2006; Frühholz et al., 2015) as well as in interoceptive processes (Engelen et al., 2023; Khalsa et al., 2018). Additionally, brain activations of the prefrontal cortex—that was also shown to be involved in interoception processing (Engelen et al., 2023; Gibson, 2019; Khalsa et al., 2018), were also revealed.

Wholebrain analysis for the Production factor alone or the interaction between Emotion and Production did not result in any above-threshold expected effects. However, we conducted a planned, mask-inclusive analysis on regions of interest, including the TVA, IFG, somatosensory and motor cortices, insula to further investigate the Production factor. This ROI analysis yielded to activations in the TVA and the IFG (*pars opercularis*, *pars orbitalis*, and *pars triangularis*). It is consistent with the literature on both the perception and production of emotional vocalizations (Guenther, & Hickok, 2015; Wright et al., 2016; Zheng et al., 2010). This result is likely to be related to the fact that our participants could hear—even though very attenuated acoustically speaking—their own voice and literature showed activations of these areas as auditory feedback and control of vocalizations (Parrell & Houde, 2019; Tourville, Reilly, & Guenther, 2008). Particularly, the TVA are involved in the provision of articulatory maps (Aziz-Zadeh et al., 2010; Pichon & Kell, 2013), in sensory motor integration (Hickok, 2009; Peschke et al., 2009), and in phonological feedback processing (Dogil et al, 2022; Parrell, & Houde, 2019). Activations within the TVA included specifically the right mid STS for normal production compared to whispering and articulating, as well as for the direct comparison between normal vs. whispering and whispering vs. articulating, emphasizing the role of this subregion of the TVA for voiced emotion production. Activations in the bilateral primary motor cortices were observed as well for voiced emotion production, especially close to the SMA in the medial portion of the brain. These regions are indeed associated with the planning of articulatory movements (Dogil et al., 2022) and the control of the motor output in speech production (Alario et al., 2006; Frühholz et al., 2015). Furthermore, activations were revealed in the left mid insula: the only significant peak in these ROI-analyses when directly comparing normal production to whispering. The insula is recruited for the execution of speech movements (Dogil et al., 2022) and has also been highlighted as a region implicated in interoceptive awareness and emotional experience (Zaki et al., 2012). It was speculated that the insular cortex could serve as a central area for the integration of somato-visceral sensations, subjective feelings, cognitive appraisals, and the consciousness of internal and external physiological signals (Adolfi et al., 2017; Casals-Gutierrez, & Abbey, 2020; Grandjean et al., 2008). This result therefore suggests the existence of key interoceptive feedback present solely in the normal production condition—the only condition involving vibratory signals, and that takes place within the left mid insula. Activations in the left primary somatosensory cortex for voiced emotion production suggests this region is—logically—engaged in interoceptive processes during normal voice production or whispering (Engelen et al., 2023; Gibson, 2019; Khalsa et al., 2018). These ROI activations align well with our research, which aimed to explore the vibratory mechanisms occurring during speech production. Our study investigated the incidental sensory processing of these vibrations—as recorded by over-the-skin throat accelerometry—originating from the vocal cords during the production of vocal emotions, with the hypothesis that these vibrations may activate the interoception network. In fact, normal voice production was the only one with a vibratory sensation, since no vibrations are emitted for whispering and articulating production conditions due to the absence of any vocal cords’ movement. In Dunn and colleagues (2010), participants had to rate emotional pictures and to complete a heartbeat detection task from which an interoceptive accuracy score was computed.

It was found that interoceptive accuracy moderates the relation between bodily responses and emotion experience, especially subjective arousal. In the same way, we hypothesized in our study that the vibrations emitted by the participants when producing vocal emotion would constitute an interoceptive signal that would influence the neural mechanisms involved in the production of emotional voice stimuli. Our results are in line with these findings and their interpretations. The *exploratory* results of the analysis of the brain correlates of vocal tract vibrations specific to normal voice production—no vibrations are emitted for whisper and articulate production conditions due to the absence of any vocal cords movement—support the previous findings even further, with activations in areas involved in interoceptive processes such as the insula, the frontal cortex (IFG, OFC, DLPFC), and different subparts of the cerebellum. The insula, as previously mentioned plays a crucial role in interoceptive processing (Adolfi et al., 2017; Casals-Gutierrez, & Abbey, 2020). For example, Haruki and Ogawa (2021) conducted a study in which participants were instructed to focus on their heartbeat and count the number of heartbeats within a given time frame. The results showed activation in the anterior insula for both interoceptive attention and accuracy. In this study, although participants were not instructed to focus on their vibrations, these can nevertheless represent an interoceptive signal processed by using similar mechanisms. Indeed, we found vibratory brain correlates in the anterior insula when comparing happy versus angry normal voice production. As for our frontal brain correlates, prefrontal cortex was previously found to be involved in the interoceptive circuitry by receiving visceral afferent information and showed activations in reaction to respiration and heartbeat as well (Berntson, & Khalsa, 2021; Engelen et al., 2023; Joyce & Barbas, 2018; Levinthal & Strick, 2020). Finally, activations in the cerebellum were associated with interoceptive awareness (Critchley et al., 2004), in the representation of interoceptive signals (Santangelo et al., 2018). A review of the literature even presented the cerebellum as “[…] an embodying machine that provides internal models to integrate bodily information and emotional responses” (Petrosini et al., 2022), and this literature is also in line with the observed brain correlates of vibrations during production in our study. Other brain correlates of vocal tract vibrations specific to normal voice production were also observed in the STG, STS and the inferior temporal sulcus. As mentioned above, they are brain areas most frequently involved in the perception of vocal emotion, but also in the production as part of auditory feedback and control of the vocal output mechanisms (Parrell & Houde, 2019; Tourville et al., 2008).

### Limits

Our study has several limitations that need to be mentioned. Firstly, we only represented three emotions: anger, happiness and neutral. We could not include a larger panel of emotions as we needed a significant number of trials per emotion, and it was not possible to design a task using more modalities without significantly losing power. We chose anger and happiness to represent positively- and negatively-valenced emotions, and neutral as a control condition. Future studies using similar paradigms should explore various other emotions. Secondly, the vibratory signal was recorded from the throat using only one accelerometer. However, other locations, such as the head (Munger & Thomson, 2008; Nolan et al., 2009) and the chest (Sundberg, 1992) are also relevant for measuring this type of vibrations. Further investigation of other emotions and recording at multiple locations would provide a more representative view of the mechanisms underlying vocal emotion production. Moreover, our vibratory correlates findings should be taken with a grain of salt, since a number of our participants were lost due to technical problems during data collection that lead to a reduced sample size of 16 individuals. We are convinced that with a larger sample, the results would have been more conclusive, especially since almost all of the expected regions of interest are present in the exploratory results. This aspect has to be carefully addressed in the future. Regarding the recording of vibrations, we analyzed the averaged amplitude of each production, and we think it may be worth considering additional acoustic parameters or advanced signal extraction from the accelerometer signal in the future in a study dedicated to the classification of specific parameters of interest. Concerning wholebrain analysis of the Production factor or its interaction with the Emotion factor, we think the absence of wholebrain results can be attributed to the experiment’s design. Indeed, the mixing of the production conditions within the blocks was probably not ideal for the participants, involving a high level of task switching. As with the Emotion factor, separate, longer blocks should have been used for each Production condition, instead of mini blocks. Indeed, production instructions may have been unsettling, and more time spent within such production block could have been beneficial for participants to become fully familiar with them. Additionally, vocal productions induce movement, requiring extremely conservative neuroimaging preprocessing steps to remove noise. Such preprocessing removed a lot of noise from the data but it may have removed signal as well, potentially linked to the production factor.

## Conclusion

In the present study, we aimed to shed new light on voice production and the associated—potential—interoceptive information it may provide the speaker with, through vocal cord vibrations concomitant to emotional speech production. Our results provide fresh evidence for a common brain network involved both in vocal emotion *perception* and *production* involving temporal, frontal, motor and somatosensory brain regions and the insular cortex that are in line with a theoretical view of the embodiment of vocal emotions. Moreover, our exploratory analysis of the brain correlates of vocal tract vibrations during normal production provides support for the contribution of sensory, interoceptive information processing to the production of vocal emotions. This work hopefully opens the way to future studies on the fascinating interactions between embodied emotions and somatosensory processing in the vocal domain, representing an important part of daily life social interactions in human society.

## Funding

This work as well as Open Access publication charges were funded by the Swiss National Science Foundation (grant number 10531C_189397/DG-LC).

## Author notes

L.C. & D.G. designed the paradigm and experimental architecture of the task, L.C. programmed the task, acquired the data and analyzed the MRI data, G.S. wrote the first draft of the manuscript, analyzed the data and edited the manuscript during the whole writing process. All authors took part in manuscript writing and editing before submission. All authors have seen and approved the manuscript, and it hasn’t been accepted or published elsewhere.

## Notes

### Competing Interest Statement

The authors have declared no competing interest.

## References

Adolfi, F., Couto, B., Richter, F., Decety, J., Lopez, J., Sigman, M., … & Ibáñez, A. (2017). Convergence of interoception, emotion, and social cognition: a twofold fMRI meta-analysis and lesion approach. Cortex, 88, 124–142.

Alario, F. X., Chainay, H., Lehericy, S., & Cohen, L. (2006). The role of the supplementary motor area (SMA) in word production. Brain research, 1076(1), 129–143.

Aziz-Zadeh L, Sheng T, Gheytanchi A. 2010. Common premotor regions for the perception and production of prosody and correlations with empathy and prosodic ability. PLoS ONE. 5. e8759.

Banse, R., & Scherer, K. R. (1996). Acoustic profiles in vocal emotion expression. Journal of personality and social psychology, 70(3), 614.

Bänziger, T., & Scherer, K. R. (2010). Introducing the geneva multimodal emotion portrayal (gemep) corpus. Blueprint for affective computing: A sourcebook, 2010, 271–94.

Berntson, G. G., & Khalsa, S. S. (2021). Neural circuits of interoception. Trends in neurosciences, 44(1), 17–28.

Brainard, D. H. (1997). The psychophysics toolbox. Spatial Vision, 10(4), 433–436. 10.1163/156856897X00357

Casals-Gutierrez, S., & Abbey, H. (2020). Interoception, mindfulness and touch: A meta-review of functional MRI studies. International Journal of Osteopathic Medicine, 35, 22–33.

Collins, D. L., Neelin, P., Peters, T. M., & Evans, A. C. (1994). Automatic 3D intersubject registration of MR volumetric data in standardized Talairach space. Journal of computer assisted tomography, 18(2), 192–205.

Craig, A. D. (2002). How do you feel? Interoception: the sense of the physiological condition of the body. Nature reviews neuroscience, 3(8), 655–666.

Critchley, H. D., Wiens, S., Rotshtein, P., Öhman, A., & Dolan, R. J. (2004). Neural systems supporting interoceptive awareness. Nature neuroscience, 7(2), 189–195.

Dogil G, Ackermann H, Grodd W, Haider H, Kamp H, Mayer J, Riecker A, Wildgruber D. 2002. The speaking brain: a tutorial introduction to fMRI experiments in the production of speech, prosody and syntax. J Neurolinguistics. 15, 59–90.

Dunn, B. D., Galton, H. C., Morgan, R., Evans, D., Oliver, C., Meyer, M., … & Dalgleish, T. (2010). Listening to your heart: How interoception shapes emotion experience and intuitive decision making. Psychological science, 21(12), 1835–1844.

Engelen, T., Solcà, M., & Tallon-Baudry, C. (2023). Interoceptive rhythms in the brain. Nature Neuroscience, 26(10), 1670–1684.

Faul, F., Erdfelder, E., Lang, A. G., & Buchner, A. (2007). G* Power 3: A flexible statistical power analysis program for the social, behavioral, and biomedical sciences. Behavior research methods, 39(2), 175–191.

Frühholz, S., Klaas, H. S., Patel, S., & Grandjean, D. (2015). Talking in fury: the cortico-subcortical network underlying angry vocalizations. Cerebral Cortex, 25(9), 2752–2762.

Gibson, J. (2019). Mindfulness, interoception, and the body: A contemporary perspective. Frontiers in psychology, 10, 2012.

Grandjean, D., Bänziger, T., & Scherer, K. R. (2006). Intonation as an interface between language and affect. Progress in brain research, 156, 235–247.

Hage, S., R. (2010). Neuronal networks involved in the generation of vocalization. In: Brudzynski SM, editor. Handbook of mammalian vocalization—an integrative neuroscience approach. Oxford: Academic Press. p. 339–349.

Haruki, Y., & Ogawa, K. (2021). Role of anatomical insular subdivisions in interoception: Interoceptive attention and accuracy have dissociable substrates. European journal of neuroscience, 53(8), 2669–2680.

Hickok, G. (2009). The functional neuroanatomy of language. Phys Life Rev. 6, 121–143.

Holland, R., Leff, A. P., Josephs, O., Galea, J. M., Desikan, M., Price, C. J., … & Crinion, J. (2011). Speech facilitation by left inferior frontal cortex stimulation. Current Biology, 21(16), 1403–1407.

Joyce, M. K. P., & Barbas, H. (2018). Cortical connections position primate area 25 as a keystone for interoception, emotion, and memory. Journal of Neuroscience, 38(7), 1677–1698.

Jurgens, U. (2009). The neural control of vocalization in mammals: a review. J Voice. 23, 1– 10.

Khalsa, S. S., Adolphs, R., Cameron, O. G., Critchley, H. D., Davenport, P. W., Feinstein, J. S., … & Zucker, N. (2018). Interoception and mental health: a roadmap. Biological psychiatry: cognitive neuroscience and neuroimaging, 3(6), 501–513.

Kleiner, M., Brainard, D., & Pelli, D. (2007). What’s new in psychtoolbox-3?

Laukka, P., Juslin, P., & Bresin, R. (2005). A dimensional approach to vocal expression of emotion. Cognition & Emotion, 19(5), 633–653.

Laukka P, Åhs F, Furmark T, Fredrikson M. 2011. Neurofunctional correlates of expressed vocal affect in social phobia. Cogn Affect Behav Neurosci. 11, 413–425.

Levinthal, D. J., & Strick, P. L. (2020). Multiple areas of the cerebral cortex influence the stomach. Proceedings of the National Academy of Sciences, 117(23), 13078–13083.

Manto, M., Bower, J. M., Conforto, A. B., Delgado-García, J. M., Da Guarda, S. N. F., Gerwig, M., … & Timmann, D. (2012). Consensus paper: roles of the cerebellum in motor control—the diversity of ideas on cerebellar involvement in movement. The Cerebellum, 11, 457–487.

Munger, J. B., & Thomson, S. L. (2008). Frequency response of the skin on the head and neck during production of selected speech sounds. The Journal of the Acoustical Society of America, 124(6), 4001–4012.

Nolan, M., Madden, B., & Burke, E. (2009). Accelerometer based measurement for the mapping of neck surface vibrations during vocalized speech. Annual International Conference of the IEEE Engineering in Medicine and Biology Society, 4453–4456.

Parkinson, A. L., Flagmeier, S. G., Manes, J. L., Larson, C. R., Rogers, B., & Robin, D. A. (2012). Understanding the neural mechanisms involved in sensory control of voice production. Neuroimage, 61(1), 314–322.

Parrell, B., & Houde, J. (2019). Modeling the role of sensory feedback in speech motor control and learning. Journal of Speech, Language, and Hearing Research, 62(8S), 2963–2985.

Pell, M. D., & Kotz, S. A. (2011). On the time course of vocal emotion recognition. PLoS One, 6(11).

Pelli, D. G. (1997). The videotoolbox software for visual psychophysics: Transforming numbers into movies. Spatial Vision, 10(4), 437–442. 10.1163/156856897X00366

Peschke, C., Ziegler, W., Kappes, J., & Baumgaertner, A. (2009). Auditory– motor integration during fast repetition: the neuronal correlates of shadowing. Neuroimage. 47, 392–402.

Petrosini, L., Picerni, E., Termine, A., Fabrizio, C., Laricchiuta, D., & Cutuli, D. (2022). The cerebellum as an embodying machine. The Neuroscientist, 10738584221120187.

Pichon, S., & Kell, C. A. (2013). Affective and sensorimotor components of emotional prosody generation. J Neurosci. 33, 1640–1650

Riecker, A., Mathiak, K., Wildgruber, D., Erb, M., Hertrich, I., Grodd, W., & Ackermann, H. (2005). fMRI reveals two distinct cerebral networks subserving speech motor control. Neurology, 64(4), 700–706.

Risberg, A., & Lubker, J. (1978). Prosody and speechreading. Speech Transmission Laboratory Quarterly Progress Report and Status Report, 4, 1–16.

Ross, E. D., & Monnot, M. (2008). Neurology of affective prosody and its functional-anatomic organization in right hemisphere. Brain Lang, 104,51–74.

Santangelo, G., Vitale, C., Baiano, C., D’Iorio, A., Longo, K., Barone, P., … & Conson, M. (2018). Interoceptive processing deficit: A behavioral marker for subtyping Parkinson’s disease. Parkinsonism & related disorders, 53, 64–69.

Sauter, D. A., & Eimer, M. (2010). Rapid detection of emotion from human vocalizations. Journal of cognitive neuroscience, 22(3), 474–481.

Scherer, K. R., & Oshinsky, J. S. (1977). Cue utilization in emotion attribution from auditory stimuli. Motivation and emotion, 1(4), 331–346.

Schulz, S. M. (2016). Neural correlates of heart-focused interoception: a functional magnetic resonance imaging meta-analysis. Philosophical Transactions of the Royal Society B: Biological Sciences, 371(1708), 20160018.

Scott, S. K. (2022). The neural control of volitional vocal production—from speech to identity, from social meaning to song. Philosophical Transactions of the Royal Society B, 377(1841), 20200395.

Sundberg, J. (1992). Phonatory vibrations in singers: A critical review. Music Perception: An Interdisciplinary Journal, 9(3), 361–381.

Tourville, J. A., Reilly, K. J., & Guenther, F. H. (2008). Neural mechanisms underlying auditory feedback control of speech. Neuroimage, 39(3), 1429–1443.

Tzourio-Mazoyer, N., Landeau, B., Papathanassiou, D., Crivello, F., Etard, O., Delcroix, N., … & Joliot, M. (2002). Automated anatomical labeling of activations in SPM using a macroscopic anatomical parcellation of the MNI MRI single-subject brain. Neuroimage, 15(1), 273–289.

Whitfield-Gabrieli, S., & Nieto-Castanon, A. (2012). Conn: a functional connectivity toolbox for correlated and anticorrelated brain networks. Brain connectivity, 2(3), 125–141.

Wright, A. E., Davis, C., Gomez, Y., Posner, J., Rorden, C., Hillis, A. E., & Tippett, D. C. (2016). Acute ischemic lesions associated with impairments in expression and recognition of affective prosody. Perspectives of the ASHA Special Interest Groups, 1(2), 82–95.

Yelnik J. (2008). Modeling the organization of the basal ganglia. Rev Neurol (Paris). 164:969– 976

Zaki, J., Davis, J. I., & Ochsner, K. N. (2012). Overlapping activity in anterior insula during interoception and emotional experience. Neuroimage, 62(1), 493–499.

Zheng, Z. Z., Munhall, K. G., & Johnsrude, I. S. (2010). Functional overlap between regions involved in speech perception and in monitoring one’s own voice during speech production. Journal of cognitive neuroscience, 22(8), 1770–1781.

