## Supplementary material for "Neural correlates of embodied and vibratory mechanisms associated with vocal emotion production"

### Supplementary information on MRI data acquisition and analysis

All functional imaging data were recorded on a 3-T Siemens Trio System (Siemens, Erlangen, Germany) scanner equipped with a 32-channel antenna. A 3D sequence was used to acquire an extremely high-resolution T1-weighted image (0.35x0.35x0.7mm; TR=2400msec; TE=2.29ms; FA=8degrees; 256 total slices).

For the voice production task, functional images were acquired in descending order using a multi-band echo-planar imaging (EPI) sequence of 56 slices aligned along the anterior-posterior commissure (voxel size: 2.5mm isotropic; slice thickness=2mm; TR=1000msec; TE=30ms; FOV=205 x 205mm; matrix resolution=64 x 64; FA=64degrees; BW=1952Hz/px). Functional image acquisition was achieved by sparse sampling to facilitate vocal output from the participants and recording by the microphone: five 1sec volumes were sampled following the vocal production period of 3sec.

For the functional temporal voice areas localizer, image acquisition was continuous and data were acquired by using a multislice echo planar imaging sequence (36 transversal slices in descending order, voxel size: 3x3x3.2mm, slice thickness=3.2mm, TR=2100msec, TE=30msec, FOV=205 x 205mm, 64 x 64 matrix, FA=90degrees, bandwidth 1562Hz/Px).

#### fMRI preprocessing and data analysis

##### Voice production task

Due to voice production in each trial and the associated head movement it inherently created, data had to be treated with high sensitivity. Preprocessing was therefore computed using a mix of SPM12 (SPM12, Wellcome Trust Centre for Neuroimaging, London, UK), the CONN toolbox (Whitfield-Gabrieli and Nieto-Castanon 2012) and the Artifact detection tools (ART) toolbox (https://www.nitrc.org/projects/artifact_detect/). Functional data were first converted to 4D Nifti in SPM12 for each run separately to get one single file per run, for more efficient computation performance. Preprocessing steps were regrouped in a custom script, including the following steps: realignment, artifact detection followed by an additional denoising (scrubbing including functional regression and functional bandpass). At this stage, normalization into the Montreal Neurological Institute (MNI) space (Collins et al. 1994) was performed for improved comparison between the brain tissues of our sample of participants. Finally, data were spatially smoothed with an isotropic Gaussian filter of 8 mm full width at half maximum. Following an initial step consisting in the creation of a participant-specific matrix including the onset and duration as well as vibratory signals of each trial of each run, preprocessed and smoothed functional images were then analyzed with SPM12 both at the participant-level (‘first-level’) as well as at the group-level (‘second-level’). Six general linear models were used to compute first-level statistics, using as trial-level covariate the Z-axis averaged vibratory signal recorded by the accelerometer located on the left side of the vocal tract on the throat, three centimeters from the glottis in the horizontal plane. All models used a convolution with a Finite Impulse Response (FIR) with the duration of the production period for each trial. Each event was therefore modelled using FIR function, locked to scans 2-3-4 (3sec duration) out of the five 1sec-volumes acquired after each trial through script-triggered sparse sampling data acquisition.

Model 1 focused on the main effect of the Emotion factor, Model 2 on the Production factor, Model 3 on the interaction between Emotion and Production factors, Model 4 on vibrations of the Emotion factor, Model 5 on the vibrations of the Production factor, Model 6 on the vibrations of the interaction between Emotion and Production factors. For each model, six motion parameters were included as regressors of no interest to account for movement in the data. For each model, regressors were used to compute simple contrasts for each participant, leading to separate main effects of: Angry, Neutral and Happy voice productions (Model 1), Normal, Whisper, Articulate and Imagine voice productions (Model 2), the interaction between these two factors for model 3 (Normal, Whisper, Articulate and Imagine for Angry, Neutral and Happy voice productions) and in a similar fashion for vibrations linked to each factor and their interaction (Model 4, 5, 6, respectively).

These models yielded to 6 flexible factorial second-level analyses: one for each model for the conditions (Emotion, Production and Emotion*Production, respectively) and one for each model for the covariate only (correlates of recorded vibrations for Emotion, Production and Emotion*Production, respectively). For all six analyses, we had the Participants factor (Factor 1 in the analysis) to take into account interindividual variability as well as the factor(s) of interest. For the Participants factor of each model, data independence was set to ‘yes’, variance to ‘unequal’ while for the factor(s) of interest data independence set to ‘no’, variance to ‘unequal’ (Model 1: Participants, Emotion as Factors 1 & 2, respectively; Model 2: Participants, Production as Factors 1 & 2, respectively; Model 3: Participants, Emotion, Production as Factors 1, 2 & 3, respectively; Model 4: Participants, Emotion vibrations as Factors 1 & 2, respectively; Model 5: Participants, Production vibrations as Factors 1 & 2, respectively; Model 6: Participants, Emotion and Production vibrations as Factors 1, 2 & 3, respectively).

Additional region-of-interest (ROI) analyses were computed for Model 3 (conditions, interaction effect of Emotion*Production), by using a mask of our ROIs, including the bilateral: pre-motor, primary motor, primary somatosensory cortices, pre-supplementary motor area, supramarginal gyrus, superior temporal cortex and temporal voice areas, inferior frontal gyrus (*pars opercularis, triangularis and orbitalis*) and the insula. The ROI mask included a total of 14'490 voxels in the left hemisphere and 13'842 voxels in the right hemisphere ($K_{total}$=28'332 voxels). These analyses were computed to investigate the interaction between Emotion and Production factors and were based on the [Anger, Happiness > Neutral] contrast thresholded in SPM12 by using a voxel-wise false discovery rate (FDR) correction at $p$<.05 and an arbitrary cluster extent of k>20, mask-inclusive. Each above-threshold peak of this contrast was used for beta extraction: for each peak activation, the nearest local maxima within a radius of 3mm was searched for each participant and then a cube of 27 contiguous voxels was used around this selected peak, computing singular value decomposition among these 27 voxels to select the ones that explained at least 90% of the variance. Statistics were then computed using repeated measures ANOVAs (Statistica v.14.1.0.8, StatSoft GmbH, Germany) with factors Emotion and Production type, for each region. Significant effects are reported in Figure 5F and all tested effects in Supplementary Table 1.

For non-ROI analyses, activations were thresholded similarly in SPM12 by using a voxel-wise FDR correction at $p$<.05 and an arbitrary cluster extent of k>20 voxels to remove small clusters of activity or single-voxel activations for Model 1, 2, 3 (N=25). For Model 4, 5, 6 (N=16) activations were thresholded in SPM12 by using a voxel-wise uncorrected $p$<.005 and an arbitrary cluster extent of k>20 voxels to remove small clusters of activity or single-voxel activations. This discrepancy is due to the exclusion of 9 participants for these latter models due to data contamination (bad accelerometer cable isolation leading to high, irreparable noise in the recorded data). Contrast activations were rendered on brains from the CONN toolbox (Whitfield-Gabrieli and Nieto-Castanon 2012). The outlines for some of our lateral regions of interest in Figures 2-3-4-5 were delineated using the latest version of the 'automated anatomical labelling' ('aal') atlas (Tzourio-Mazoyer et al. 2002) implemented in the 'WFU Pickatlas' toolbox (https://www.nitrc.org/projects/wfu_pickatlas).

**Temporal voice areas functional localizer**

Functional images were analyzed with Statistical Parametric Mapping software (SPM12, Wellcome Trust Centre for Neuroimaging, London, UK). Preprocessing steps included realignment to the first volume of the time series, slice timing, normalization into the

Montreal Neurological Institute (MNI) space and spatial smoothing with an isotropic Gaussian filter of 8 mm full width at half maximum.

A general linear model was used to compute first-level statistics, in which each block was modeled by using a block function and was convolved with the hemodynamic response function, time-locked to the onset of each block. Separate regressors were created for each condition (vocal and non-vocal; Condition factor). Finally, six motion parameters were included as regressors of no interest to account for movement in the data. The condition regressors were used to compute simple contrasts for each participant, leading to a main effect of vocal and non-vocal at the first-level of analysis ([1 0] for vocal, [0 1] for non-vocal). These simple contrasts were then taken to a flexible factorial second-level analysis in which there were two factors: the Participants factor (independence set to 'yes', variance set to 'unequal') and the Condition factor (independence set to 'no', variance set to 'unequal'). All neuroimaging activations were thresholded in SPM12 by using a voxel-wise FDR correction at $p<.05$ and an arbitrary cluster extent of $k>20$ voxels. These data are represented by a white outline named 'TVA'—for "Temporal Voice Areas"—in Figure 2-3-4 and by a black outline in Figure 5-6, with the original thresholded contrast images of this task reported in Figure S1.

### Supplementary figures

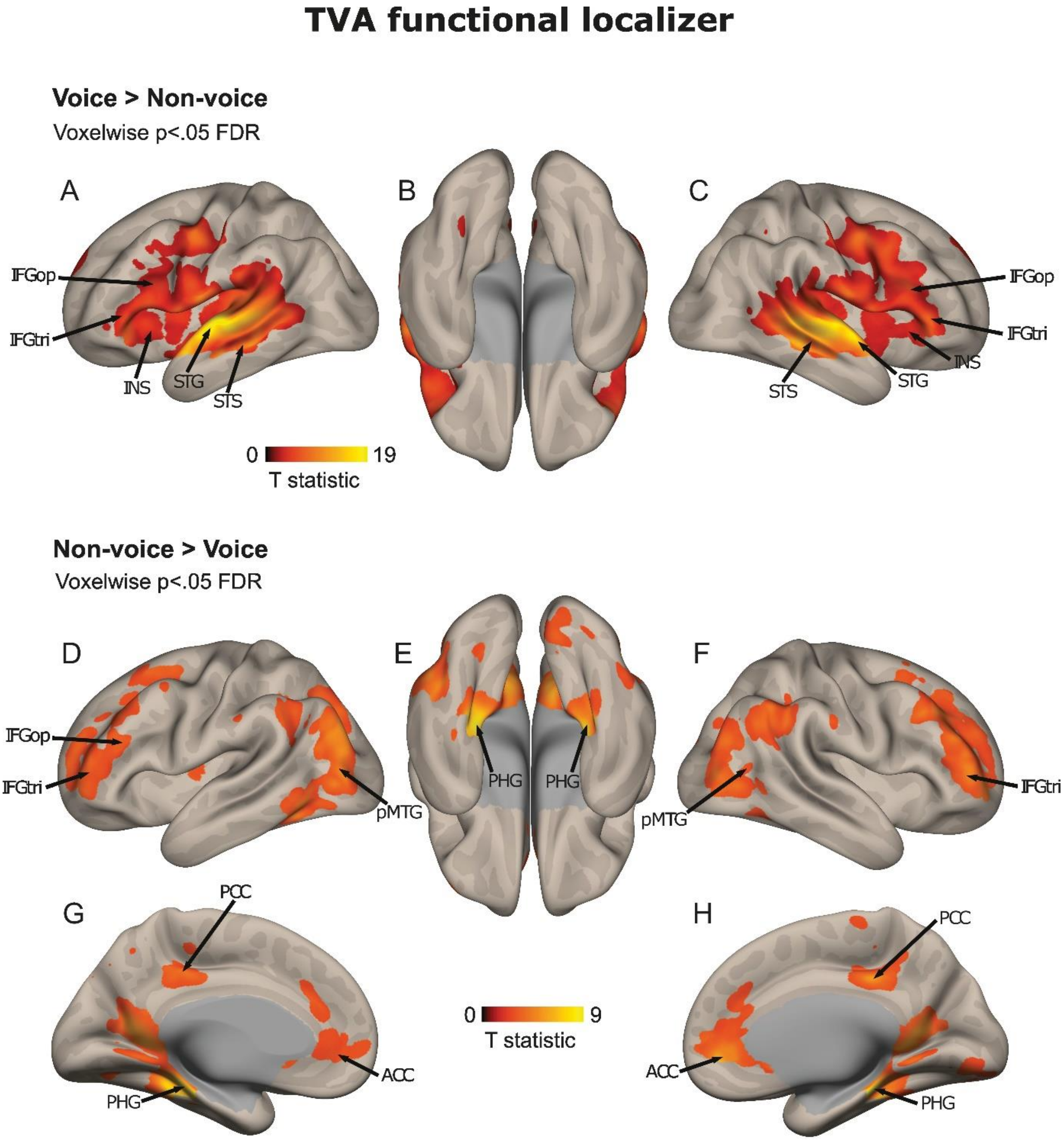


*Supplementary figure 1.* Temporal voice areas functional localizer (wholebrain voxel-wise $p < .05$ FDR, k > 20 voxels). The colorbars represents the statistical T value. ACC: anterior cingulate cortex, IFG: inferior frontal gyrus, INS: insula, op: pars opercularis, orb: pars orbitalis, PCC: posterior cingulate cortex, PHG: parahippocampal gyrus, pMTG: posterior middle temporal gyrus, STG: superior temporal gyrus, STS: superior temporal sulcus, tri: pars triangularis.

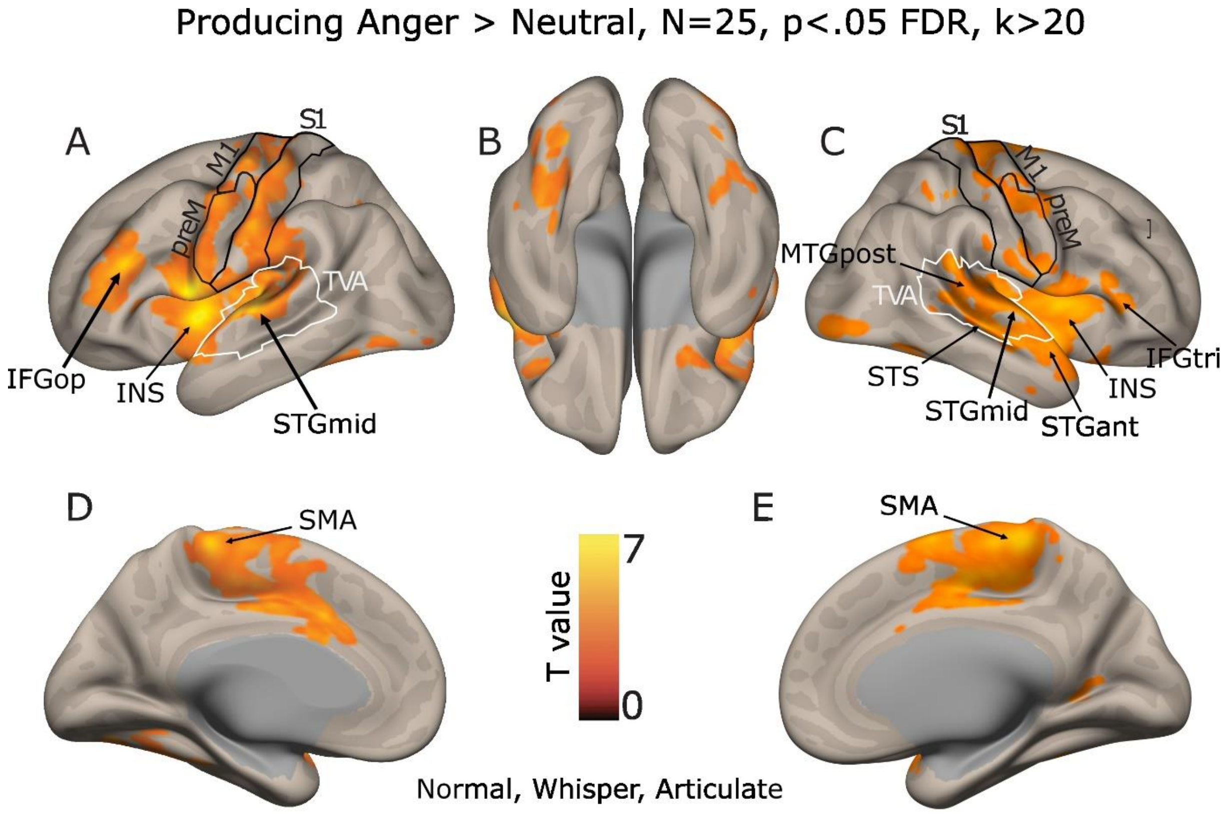


*Supplementary Figure 2.* Brain measures for the production of angry compared to neutral voices. Statistical thresholding was set to an p<.05 FDR, k>20. The colorbar represents the statistical T value. IFG: inferior frontal gyrus ('tri': *pars triangularis*, 'op': *pars opercularis*), M1: primary motor cortex, preM: premotor area, S1: primary somatosensory cortex, SMA: supplementary motor area, STGmid/ant: middle/anterior superior temporal gyrus, INS: insula, MTGpost: posterior middle temporal gyrus, STS: superior temporal sulcus, TVA: temporal voice areas delineated by white line shapes.

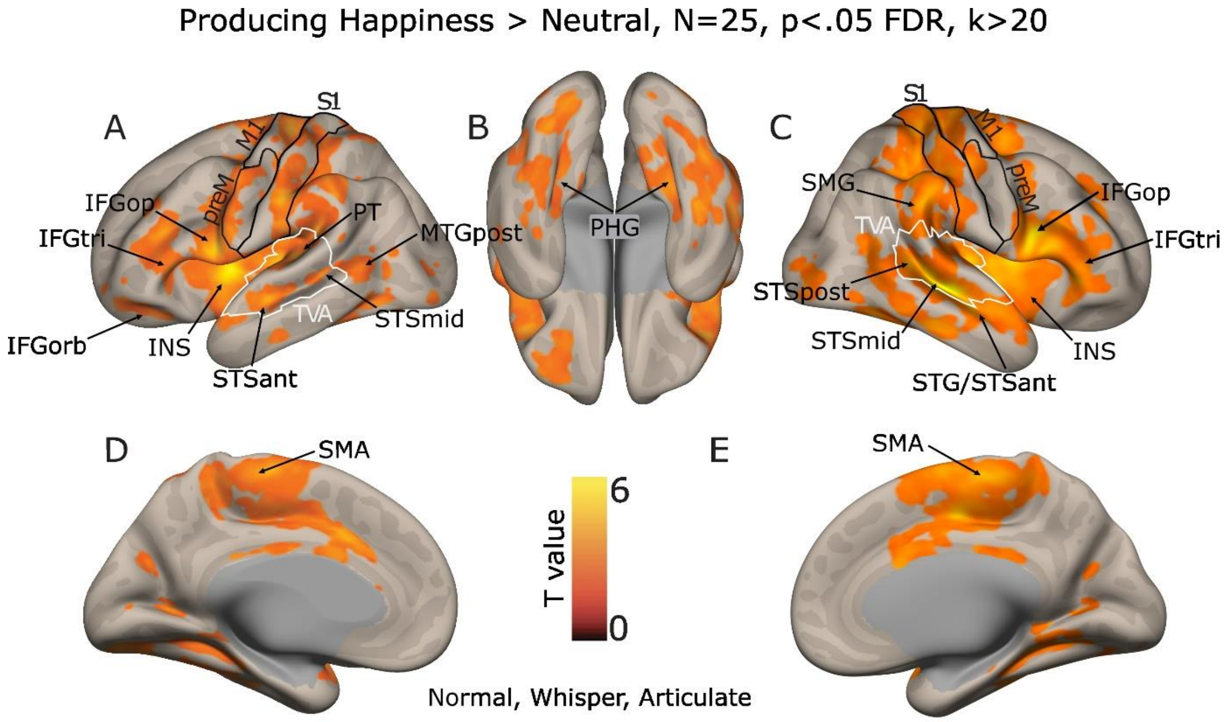


*Supplementary Figure 3.* Brain measures for the production of happy compared to neutral voices. Statistical thresholding was set to an p<.05 FDR, k>20. The colorbar represents the statistical T value. IFG: inferior frontal gyrus ('tri': *pars triangularis*, 'op': *pars opercularis*, 'orb': *pars orbitalis*), M1: primary motor cortex, preM: premotor area, S1: primary somatosensory cortex, SMA: supplementary motor area, STGmid/ant: middle/anterior superior temporal gyrus, INS: insula, MTGpost: posterior middle temporal gyrus, SMG: supramarginal gyrus, PT: planum temporale, STSmid/post: mid/posterior superior temporal sulcus, TVA: temporal voice areas delineated by white line shapes.

**Correlates of vocal tract vibrations, production factor (N=16, p<.005 unc., k>20)**

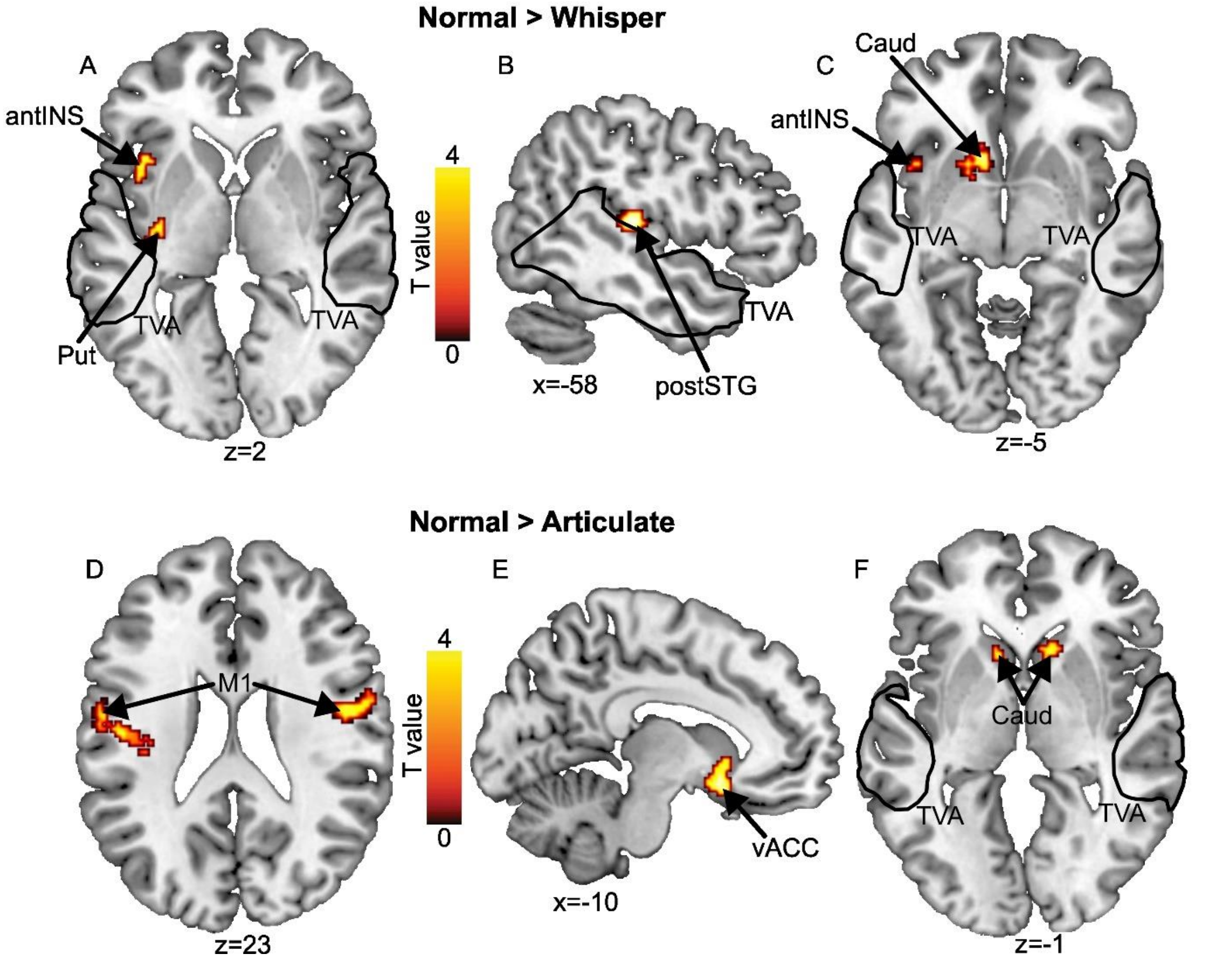


*Supplementary Figure 4.* Correlates of vocal tract vibrations for A, B and C) ‘Normal’ compared to ‘Whisper’ production condition, D, E and F) ‘Normal’ compared to ‘Articulate’ production condition. Statistical thresholding was set to an uncorrected wholebrain voxel-wise $p < .005$, $k > 20$ voxels. The colorbars represent the statistical T value. antINS: anterior insula, Caud: caudate nucleus, FP: frontal pole, IPS: intraparietal sulcus, M1: primary motor cortex, postSTG: posterior superior temporal gyrus, postSTS: posterior part of the superior temporal sulcus, Put: putamen, TVA: temporal voice areas delineated by black line shapes, vACC: ventral part of the anterior cingulate cortex.

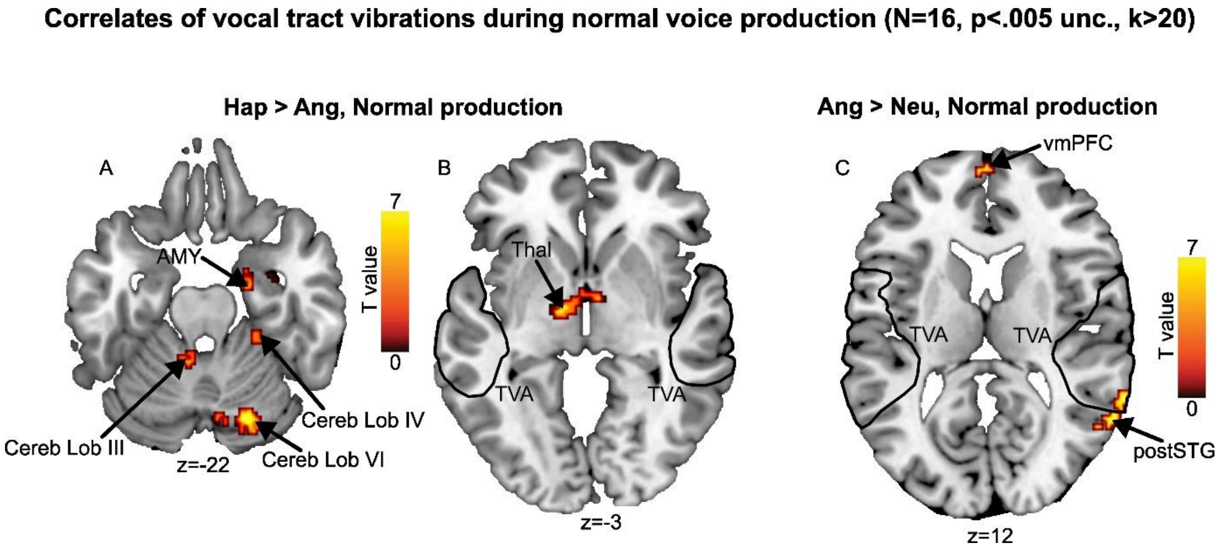


*Supplementary Figure 5.* Brain measures for the production of A and B) joyful compared to angry voices, and C) angry compared to neutral voices. Statistical thresholding was set to an uncorrected wholebrain voxel-wise $p < .005$, $k > 20$ voxels. The colorbars represents the statistical T value. AMY: amygdala, Cereb: cerebellum, Lob: lobule, postSTG: posterior part of the superior temporal gyrus, Thal: thalamus, TVA: temporal voice areas delineated by black line shapes, vmPFC: ventromedial prefrontal cortex.

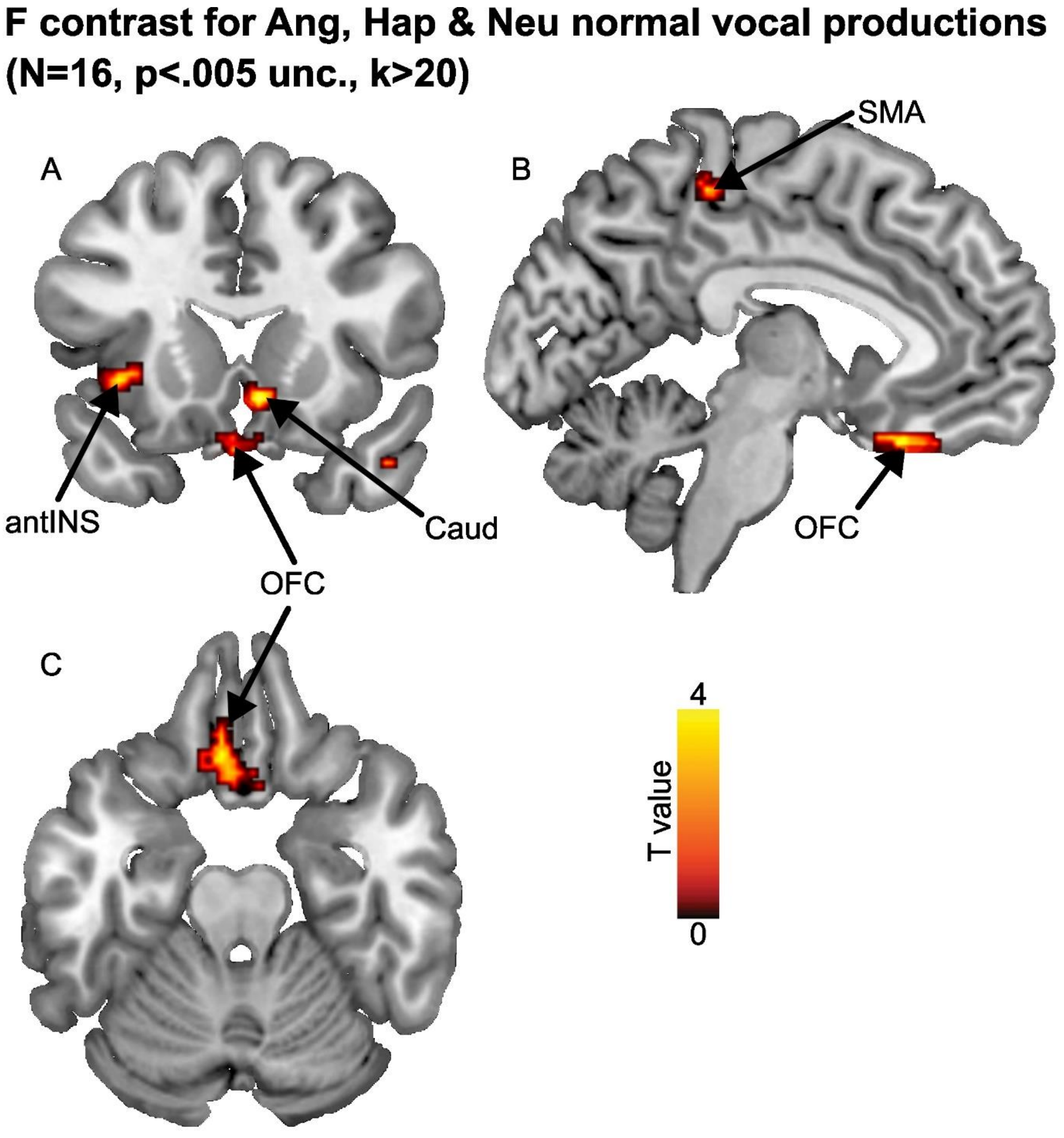


*Supplementary Figure 6.* Correlates of vocal tract vibrations for all conditions taken together (angry, happy and neutral normal productions). Statistical thresholding was set to an uncorrected wholebrain voxel-wise $p < .005$, $k > 20$ voxels. The colorbar represent the statistical T value. antINS: anterior insula, Caud: caudate nucleus, SMA: supplementary motor area, OFC: orbitofrontal cortex.

### Supplementary tables

***Supplementary Table 1.*** **Above-threshold brain clusters for the Emotion factor, angry, happy > neutral voice production (voxelwise p<.05 FDR, k>20).**

| Hemisphere | Region | MNI *X* | MNI *Y* | MNI *Z* | T-value | Cluster size |
|---|---|---|---|---|---|---|
| *L* | *mid insula* | *-38* | *4* | *2* | *10.29* | *35290* |
| R | mid STS | 44 | -22 | -4 | 9.07 | |
| R | M1 | 6 | -30 | 66 | 8.93 | |
| R | posterior STG | 48 | -24 | -2 | 8.30 | |
| L | anterior insula | -46 | 8 | 6 | 8.28 | |
| R | mid insula | 32 | 4 | 10 | 8.19 | |
| L | IFG opercularis | -52 | 6 | 6 | 8.05 | |
| R | IFG triangularis | 44 | 28 | 2 | 7.74 | |
| L | M1 | -50 | -24 | 46 | 7.71 | |
| *L* | *posterior MTG* | *-56* | *-56* | *6* | *6.13* | *187* |
| *L* | *OFC* | *-26* | *34* | *-14* | *5.62* | *114* |
| *R* | *DLPFC* | *26* | *44* | *36* | *4.13* | *117* |

STS: superior temporal sulcus; M1: primary motor cortex; STG: superior temporal gyrus; IFG: inferior frontal gyrus; MTG: middle temporal gyrus; OFC: orbitofrontal cortex; DLPFC: dorsolateral prefrontal cortex; L: left; R: right.

***Supplementary Table 2.* Above-threshold brain clusters for the Emotion factor, angry > happy voice production (voxelwise p<.05 FDR, k>20).**

| **Hemisphere** | **Region** | **MNI *X*** | **MNI *Y*** | **MNI *Z*** | **T-value** | **Cluster size** |
|---|---|---|---|---|---|---|
| *L* | *mid STG* | *-64* | *-16* | *4* | *8.29* | *731* |
| L | anterior STG | -58 | 4 | 2 | 6.91 | |
| L | posterior STG | -66 | -24 | 10 | 5.56 | |
| *R* | *Cerebellum Vermis V* | *2* | *-68* | *-10* | *7.35* | *970* |
| L | Cereb. Crus I | -32 | -74 | -22 | 6.58 | |
| L | Cereb. lobule VI | -30 | -54 | -22 | 6.24 | |
| L | vmPFC | -6 | 40 | -22 | 7.30 | |
| *C* | *S1* | *0* | *-36* | *68* | *6.68* | *937* |
| R | preSMA | 2 | 8 | 74 | 5.78 | |
| R | M1 | 8 | -18 | 80 | 5.61 | |
| *R* | *Cereb. Crus I* | *26* | *-86* | *-26* | *6.30* | *860* |
| R | Cereb. Crus I | 52 | -62 | -28 | 6.02 | |
| R | Cereb. Lobule VI | 30 | -66 | -22 | 5.10 | |
| *R* | *mid STG* | *66* | *-12* | *8* | *6.29* | *353* |
| R | anterior STG | 62 | 0 | 0 | 6.08 | |
| R | IFG opercularis | 64 | -2 | 16 | 4.72 | |
| *L* | *M1* | *-50* | *-14* | *56* | *5.68* | *225* |
| *L* | *DLPFC* | *-54* | *22* | *28* | *5.65* | *39* |
| *L* | *Thalamus* | *-10* | *-34* | *0* | *5.57* | *53* |
| *L* | *DLPFC* | *-24* | *60* | *22* | *5.34* | *107* |
| *R* | *M1* | *56* | *0* | *48* | *5.34* | *115* |
| *R* | *Amygdala* | *26* | *2* | *-14* | *4.86* | *157* |
| R | anterior insula | 44 | 4 | -12 | 3.99 | |
| *R* | *vmPFC* | *6* | *58* | *0* | *4.57* | *57* |
| *L* | *M1* | *-60* | *-2* | *36* | *4.42* | *31* |
| *L* | *S1* | *-48* | *-34* | *60* | *4.34* | *45* |

STG: superior temporal gyrus; Cereb.: cerebellar; Crus: anterior portion of the cerebral peduncle; vmPFC: ventromedial prefrontal cortex; S1: primary somatosensory cortex; preSMA: pre supplementary motor area; M1: primary motor cortex; IFG: inferior frontal gyrus; DLPFC: dorsolateral prefrontal cortex; L: left; R: right; C: central.

***Supplementary Table 3.*** **Above-threshold brain clusters for the Emotion factor, happy > angry voice production (voxelwise p<.05 FDR, k>20).**

| Hemisphere | Region | MNI *X* | MNI *Y* | MNI *Z* | T-value | Cluster size |
|---|---|---|---|---|---|---|
| *L* | *posterior MTG* | *-38* | *-58* | *12* | *10.18* | *58807* |
| R | posterior CG | 12 | -30 | 32 | 9.23 | |
| L | Hippocampus | -28 | -28 | -14 | 9.04 | |
| R | posterior MTG | 42 | -30 | -4 | 8.72 | |
| L | posterior CG | -16 | -26 | 34 | 8.69 | |
| L | Cereb. lobule V | -12 | -46 | -24 | 8.35 | |
| R | DLPFC | 26 | 22 | 28 | 8.29 | |
| R | IFG opercularis | 48 | 12 | 12 | 7.48 | |
| L | brainstem | -8 | -12 | -6 | 7.34 | |
| L | posterior STS | -54 | -34 | -4 | 6.90 | |
| L | IFG opercularis | -46 | 20 | 12 | 6.87 | |
| L | anterior STG | -52 | -6 | -8 | 6.86 | |
| R | SMA | 16 | -12 | 58 | 6.79 | |
| R | S1 | 54 | -28 | 40 | 6.39 | |
| R | mid STG | 64 | -14 | -6 | 6.27 | |
| R | Cereb. lobule V | 20 | -48 | -30 | 6.26 | |
| R | mid STS | 62 | -16 | -8 | 6.24 | |
| R | posterior STS | 56 | -40 | 2 | 6.16 | |
| R | IFG triangularis | 44 | 26 | -10 | 5.94 | |
| R | anterior insula | 38 | 12 | -18 | 5.47 | |
| L | IFG orbitalis | -44 | 42 | -16 | 5.47 | |
| L | M1 | -10 | -20 | 60 | 5.24 | |
| R | parahippocampal gyrus | 24 | -16 | -28 | 5.12 | |
| R | vmPFC | 22 | 22 | -14 | 5.00 | |
| L | IFG triangularis | -52 | 20 | 6 | 4.90 | |
| R | posterior insula | 38 | -10 | 10 | 4.58 | |
| L | amygdala | -16 | -4 | -20 | 3.70 | |

MTG: middle temporal gyrus; CG: cingulate gyrus; Cereb.: cerebellar; DLPFC: dorsolateral prefrontal cortex; IFG: inferior frontal gyrus; STG: superior temporal gyrus; STS: superior temporal sulcus; vmPFC: ventromedial prefrontal cortex; S1: primary somatosensory cortex; SMA: supplementary motor area; M1: primary motor cortex; L: left; R: right.

***Supplementary Table 4.*** **Above-threshold brain clusters for ROI-analyses (mask-inclusive) for the Production factor, corrected for multiple comparisons (voxelwise $p < .05$ FDR, k > 20 voxels) based on the [Anger, Happiness > Neutral] Emotion contrast.**

| **Hemisphere** | **Region** | **MNI *X*** | **MNI *Y*** | **MNI *Z*** | **T-value** | **Cluster size** |
|---|---|---|---|---|---|---|
| *L* | *mid insula* | *-38* | *4* | *2* | *10.00* | *2551* |
| L | posterior insula | -42 | -24 | 8 | 7.70 | |
| L | anterior insula | -44 | 4 | -6 | 7.00 | |
| *R* | *posterior STG* | *44* | *-22* | *-4* | *9.08* | *3288* |
| R | IFG triangularis | 44 | 28 | 2 | 7.51 | |
| R | posterior insula | 36 | -14 | 4 | 7.43 | |
| R | mid insula | 36 | 6 | 4 | 7.35 | |
| *R* | *S1* | *6* | *-30* | *66* | *8.87* | *644* |
| R | M1 | 4 | -22 | 66 | 7.44 | |
| R | SMA | 6 | -12 | 64 | 6.78 | |
| R | preSMA | 6 | 0 | 66 | 6.11 | |
| *L* | *S1* | *-50* | *-24* | *46* | *7.76* | *978* |
| L | preM1 | -56 | -18 | 32 | 6.84 | |
| L | SMA | -52 | -2 | 48 | 6.03 | |
| *L* | *M1* | *-2* | *-24* | *64* | *7.15* | *370* |
| L | SMA | -6 | -16 | 66 | 7.14 | |
| L | S1 | -2 | -34 | 64 | 5.37 | |
| *R* | *M1* | *34* | *-20* | *54* | *5.52* | *268* |
| R | S1 | 46 | -24 | 62 | 5.19 | |
| R | SMA | 40 | -4 | 48 | 4.92 | |
| *R* | *S1* | *18* | *-28* | *76* | *5.32* | *179* |
| *R* | *M1* | *48* | *-16* | *46* | *4.31* | *92* |
| *L* | *posterior STS* | *-56* | *-50* | *6* | *4.18* | *37* |
| *R* | *preSMA* | *4* | *12* | *58* | *3.92* | *24* |

STG: superior temporal gyrus; IFG: inferior frontal gyrus; S1: primary somatosensory cortex; M1: primary motor cortex; SMA: supplementary motor area; preSMA: pre supplementary motor area; STS: superior temporal sulcus; L: left; R: right.

***Supplementary Table 5.*** **P-values from post-hoc contrasts computed with Tukey's HSD (honestly significant difference) test in the right mid STS.**

| Cell number | Emotion | Production | 2 | 3 | 4 | 5 | 6 | 7 | 8 | 9 |
|---|---|---|---|---|---|---|---|---|---|---|
| 1 | Anger | Normal | n.s. | <.01 | n.s. | n.s. | n.s. | n.s. | <.05 | <.001 |
| 2 | Anger | Whisper | | <.01 | n.s. | n.s. | n.s. | n.s. | <.05 | <.001 |
| 3 | Anger | Articulate | | | <.001 | <.001 | <.001 | n.s. | n.s. | n.s. |
| 4 | Happiness | Normal | | | | n.s. | n.s. | <.001 | <.001 | <.001 |
| 5 | Happiness | Whisper | | | | | n.s. | <.01 | <.001 | <.001 |
| 6 | Happiness | Articulate | | | | | | <.001 | <.001 | <.001 |
| 7 | Neutral | Normal | | | | | | | n.s. | <.01 |
| 8 | Neutral | Whisper | | | | | | | | <.05 |
| 9 | Neutral | Articulate | | | | | | | | |

***Supplementary Table 6.*** **P-values from post-hoc contrasts computed with Tukey's HSD (honestly significant difference) test in the left mid insula.**

| Cell number | Emotion | Production | 2 | 3 | 4 | 5 | 6 | 7 | 8 | 9 |
|---|---|---|---|---|---|---|---|---|---|---|
| 1 | Anger | Normal | n.s. | n.s. | n.s. | n.s. | n.s. | <.001 | <.001 | <.001 |
| 2 | Anger | Whisper | | <.05 | n.s. | n.s. | <.01 | <.05 | <.01 | <.001 |
| 3 | Anger | Articulate | | | n.s. | <.001 | n.s. | <.001 | <.001 | <.001 |
| 4 | Happiness | Normal | | | | n.s. | <.05 | <.01 | <.001 | <.001 |
| 5 | Happiness | Whisper | | | | | <.001 | n.s. | n.s. | <.01 |
| 6 | Happiness | Articulate | | | | | | <.001 | <.001 | <.001 |
| 7 | Neutral | Normal | | | | | | | n.s. | n.s. |
| 8 | Neutral | Whisper | | | | | | | | n.s. |
| 9 | Neutral | Articulate | | | | | | | | |

***Supplementary Table 7.*** **Exploratory above-threshold (p<.005 uncorrected, k>20) brain clusters correlating with surface throat vibratory signals during voice production, for the Production factor separately and for its interaction with Emotion.**

| **Hemisphere** | **Region** | **MNI *X*** | **MNI *Y*** | **MNI *Z*** | **T-value** | **Cluster size** |
|---|---|---|---|---|---|---|
| Normal production > Whispering, Articulating | | | | | | |
| *L* | *posterior STG*** | *-48* | *-26* | *12* | *3.47* | *72* |
| *R* | *mid STS** | *66* | *-32* | *-4* | *3.27* | *36* |
| *L* | *SMG** | *-54* | *-40* | *48* | *3.23* | *50* |
| *L* | *Caudate nucleus** | *-10* | *16* | *-4* | *3.17* | *37* |
| L | Nucleus accumbens | -12 | 14 | -10 | 3.02 | |
| *L* | *DLPFC** | *-40* | *46* | *10* | *3.13* | *51* |
| Anger, Happy > Neutral * Normal > Whispering, Articulating production | | | | | | |
| *L* | *OFC*** | *-6* | *26* | *-22* | *3.92* | *293* |
| *R* | *posterior MTG*** | *44* | *-54* | *14* | *3.89* | *178* |
| *L* | *Cerebellum Crus I*** | *-10* | *-74* | *-20* | *3.66* | *133* |
| *R* | *SMA*** | *6* | *-18* | *58* | *3.35* | *28* |
| *R* | *anterior ITS** | *50* | *-6* | *-28* | *3.25* | *33* |
| Anger > Happy * Normal > Whispering, Articulating production | | | | | | |
| *L* | *vmPFC*** | *-12* | *52* | *-2* | *4.09* | *138* |
| *L* | *anterior CG*** | *-14* | *40* | *26* | *3.64* | *113* |
| *L* | *medial PFC*** | *-6* | *56* | *38* | *3.62* | *139* |
| *R* | *OFC*** | *2* | *18* | *-24* | *3.54* | *55* |
| *L* | *posterior STG*** | *-50* | *-44* | *20* | *3.53* | *34* |
| *R* | *anterior MTG*** | *44* | *12* | *-30* | *3.49* | *51* |
| *L* | *Cereb. lobule VI*** | *-24* | *-66* | *-26* | *3.37* | *30* |
| *L* | *posterior MTG** | *-46* | *-52* | *4* | *3.26* | *46* |
| Happy > Anger * Normal > Whispering, Articulating production | | | | | | |
| *L* | *IFG opercularis*** | *-60* | *10* | *10* | *3.72* | *21* |
| *L* | *anterior insula** | *-38* | *16* | *-4* | *3.26* | *38* |
| *R* | *fusiform cortex** | *20* | *-50* | *-8* | *3.18* | *61* |

STG: superior temporal gyrus; STS: superior temporal sulcus; SMG: supramarginal gyrus; DLPFC: dorsolateral prefrontal cortex; OFC: orbitofrontal cortex; MTG: middle temporal gyrus; Cereb.: cerebellar; Crus: anterior portion of the cerebral peduncle; SMA: supplementary motor area; ITS: inferior temporal sulcus; vmPFC: ventromedial prefrontal cortex; CG: cingulate gyrus; PFC: prefrontal cortex; IFG: inferior frontal gyrus; L: left; R: right. * survives thresholding at p<.001 uncorrected; ** survives thresholding at p<.0001 uncorrected.
